# Chromosomal instability and chromosome 17p loss drive convergent *NDE1* synthetic lethality in metastatic cancer cells

**DOI:** 10.64898/2026.02.10.704921

**Authors:** Caroline M. Teddy, Jacob P. Hoj, Julia Caci, Min Lu, Shane T. Killarney, Tal Ben-Yishay, Uri Ben-David, Kris C. Wood

## Abstract

Recent studies have identified recurrent features of metastatic cancer cells such as their increased chromosomal instability (CIN) and frequent loss of the short arm of chromosome 17 (Chr17p). However, it remains unclear whether these features induce synthetic lethal vulnerabilities that can be used to specifically target metastatic disease. Using whole-genome CRISPR/Cas9 loss-of-function screens performed in matched primary and CIN-high brain-metastatic tumor models, we discovered that brain-metastatic cells exhibit increased sensitivity to the loss of diverse regulators of chromosome segregation. Knockout of one such regulator, *NDE1,* selectively inhibited the growth of brain-metastatic models *in vitro* and *in vivo*, an effect driven by the loss of STAG2 and consequent induction of CIN. Surprisingly, dependence on *NDE1* was also highly correlated with loss of Chr17p across hundreds of cancer cell lines in DepMap, the result of losing the *NDE1* paralog *NDEL1*, which resides at this locus. CIN and Chr17p loss are thus independently sufficient to drive *NDE1* dependence in brain-metastatic cells, and the presence of both features increases *NDE1* dependence additively. These findings demonstrate that metastasis evolution endows cancer cells with specific vulnerabilities, including one that is driven by two recurrently altered molecular features of metastatic disease.

## Background

Cancer progression requires tumor cell adaptation to immense stress imposed by the changing cellular environment. This cellular stress, driven by factors that include oncogenic signaling, altered immune and metabolic environments, and drug treatments, exerts evolutionary pressures on a tumor cell population, requiring the population to make fitness trade-offs to survive. Fitness trade-offs in the context of drug treatment have been well-established by our group and others, whereby tumor cells that have adapted to drug treatment through the acquisition of various resistance mechanisms are also endowed with concomitant cellular vulnerabilities as a result of their adaptation [1–4]. These vulnerabilities, referred to as ‘collateral sensitivities’, provide attractive avenues for therapeutic development, because they can serve as the basis for evolutionary traps — by undergoing the cellular evolution required for survival, these associated vulnerabilities are also acquired. Beyond drug treatment, there are many other instances in which intense selection pressures are exerted on cancer cells, driving disease evolution. One such instance is metastasis, during which a tumor cell population is exposed to massive cycles of selection and elimination before acquiring the necessary features to spread from the primary site. Because metastatic tumor cells must acquire specific adaptations that allow for their survival, colonization, and eventual expansion at distal organ sites, we hypothesized that they must also develop cell-intrinsic collateral vulnerabilities that are distinct from the primary tumor cells from which they were derived.

Previous large-scale sequencing studies comparing primary and metastatic tumors have revealed few shared actionable mutations consistent among patients or sites of metastasis. In one of the largest sequencing cohorts, Memorial Sloan Ketting - Metastatic Events and Tropisms (MSK-Met, n=25,000), consistent increases in the frequency of mutations of a few genes, including *TP53*, *HER2* and *TERT*, were observed in metastatic tumor tissue compared to non-metastatic tissue [5]. However, of the mutations identified, few were shared in more than one tumor type or metastatic site. Conversely, MSK-Met identified chromosomal instability (CIN) as a highly recurrent feature of metastasis, with the fraction of the genome altered (FGA) significantly higher in metastatic samples in 32% of tumor types [5]. Subsequent studies have consistently identified CIN as a robust, metastasis-enriched feature that is thought to function by increasing the evolvability of tumor cell populations [6] and inducing pro-metastatic inflammatory signaling [7, 8]. High-resolution single-cell sequencing studies of brain-metastatic patient samples have identified a CIN-high subpopulation of cells which is shared across patients and particularly aggressive [9]. Given this, we sought to identify acquired vulnerabilities of metastatic cells exhibiting increased CIN, mirroring the human disease.

We identified acquired genetic vulnerabilities of metastatic cells by performing a whole-genome CRISPR/Cas9 screen comparing isogenic lung cancer cells with matched, CIN-high brain-metastatic derivatives, evolved by serial *in vivo* selection to acquire potent metastatic capacity. Several of the genes identified in the screen encoded proteins that regulate centrosome function and chromosome segregation, implicating these processes as ‘dependency hubs’ in metastatic cell lines. One acquired vulnerability of brain-metastatic cells identified by our screen, an increased dependence on NudE Neurodevelopment Gene 1 (*NDE1*), is driven by CIN and validated as a conserved vulnerability in a panel of diverse brain-metastatic lung cancer cell lines relative to their matched parental counterparts. Surprisingly, mechanistic studies revealed that dependency on *NDE1* can also be derived from the heterozygous loss of the short arm of chromosome 17 (Chr17p) – a recurrent event in certain metastatic tumors – via the loss of *NDE1*’s Chr17p-resident paralog, *NDEL1* [5, 10–13]. Subsequent experiments revealed that while Chr17p loss and CIN are linked by *TP53*, whose loss on Chr17p can facilitate CIN and metastasis [7, 8, 10, 12], they are nevertheless independently sufficient to induce *NDE1* synthetic lethality in cancer cells. This effect is strengthened when both features were present, a common occurrence in patients with brain metastasis from the lung [5]. Thus, genetic dependence on *NDE1* in brain-metastatic cancer cells represents, to our knowledge, one of the first examples of a cell-autonomous dependency enriched by metastasis evolution. Further, this phenomenon is a unique example of a ‘convergent’ synthetic lethality, in which tumor cell evolution results in two traits that are beneficial to a cancer cell (in this case, by facilitating brain metastasis) but also result in exquisite dependence on the same gene.

## Results

### CIN-high metastatic cells are dependent on mitotic regulators

To define acquired vulnerabilities of brain-metastatic cancer cells, we created isogenic parental-metastatic cell line pairs suitable for comparative genetic screening. Brain-metastatic derivatives of lung adenocarcinoma cell lines (PC9, H2030, and HCC4006) were created via serial intracardiac injection and subsequent harvest of metastatic brain lesions from mice as previously described [14, 15]. The resulting cells were brain-tropic when injected into mice and expressed transcriptional signatures associated with neuronal progenitors, consistent with other studies using similar models (Fig. S1a-b) [16, 17]. Brain-metastatic derivative cells uniformly displayed evidence of increased chromosomal instability when compared to parental cell lines, including significant increases in cGAS+ micronuclei (Fig. 1a, S1c) and centrosome duplication (Fig. 1b, S1d). Low-pass whole genome sequencing (LP-WGS) identified some additional large-scale and arm-level genomic alterations in BrM cells compared to matched parental cells, consistent with an underlying inability to faithfully segregate chromosomes (Fig. S1e). Interestingly, newly acquired aneuploidies were generally inconsistent across BrM models derived from different cell lines, suggesting that aneuploidies resulting from CIN may be random or context-dependent, similar to other published models of CIN [18, 19]. Given that BrM models maintain their increased metastatic capacity and brain tropism even after *in vitro* culture for several months, we hypothesized that vulnerabilities unique to the brain-metastatic cell state could be identified through *in vitro* genetic screens. We performed a genome-wide 3D spheroid-based CRISPR/Cas9 screen using one of our brain metastasis models (PC9-BrM3) and its matched parental cell line (PC9) (Fig. 1c). Genes whose loss is known to cause growth defects (e.g., *EGFR*) or growth advantages (e.g., *PTEN*) in PC9 cells were similarly resolved between PC9 and PC9-BrM3 cells, indicating screen fidelity (Fig. S2a-b). While, initially, a 3D screening format was chosen in hopes of better recapitulating tumor growth *in vivo* [20], screen results were highly concordant with 2D screens performed in PC9 cells in DepMap, and downstream analyses were therefore conducted in 2D culture conditions (Fig. S2c). PC9 and PC9-BrM3 cells had similar growth rates throughout the screen (Fig. S2d). Though most gene knockouts had similar effects on PC9 and PC9-BrM3 cells, the screen identified sets of genes more essential for the survival of brain-metastatic lung cancer cells than their matched parental counterparts (PC9-BrM3-biased dependencies), and genes with the reciprocal effect, which were more essential in parental cell lines compared to brain-metastatic derivatives (Parental-biased) (Fig. 1d). Known dependencies of CIN-high, aneuploid cell lines emerged as BrM3-biased genes, including *BUB1B* and *MAD2L1* (Fig. S2e) [21]. Subsequent downstream Cytoscape analysis (Fig. 1e) and Gene Set Enrichment Analysis (GSEA) (Fig. 1f) of BrM3-biased fitness genes revealed a significant overrepresentation of genes involved in mitotic regulation. In particular, PC9-BrM3 lines were strongly and differentially dependent on genes that regulate the centrosome and microtubules (Fig. 1g, Fig. S2f), implicating these processes as potential ‘dependency hubs’ in brain-metastatic cells. Consistent with these findings, *KIF2A* and *NEK1*, genes encoding mitotic regulators of centrosome function, scored as metastasis-selective dependencies that we validated *in vitro* and *in vivo* (Fig. S3). Together, these results demonstrate that CIN-high metastatic cells acquire an increased dependency on genes involved in mitotic regulation.

**Fig. 1.**
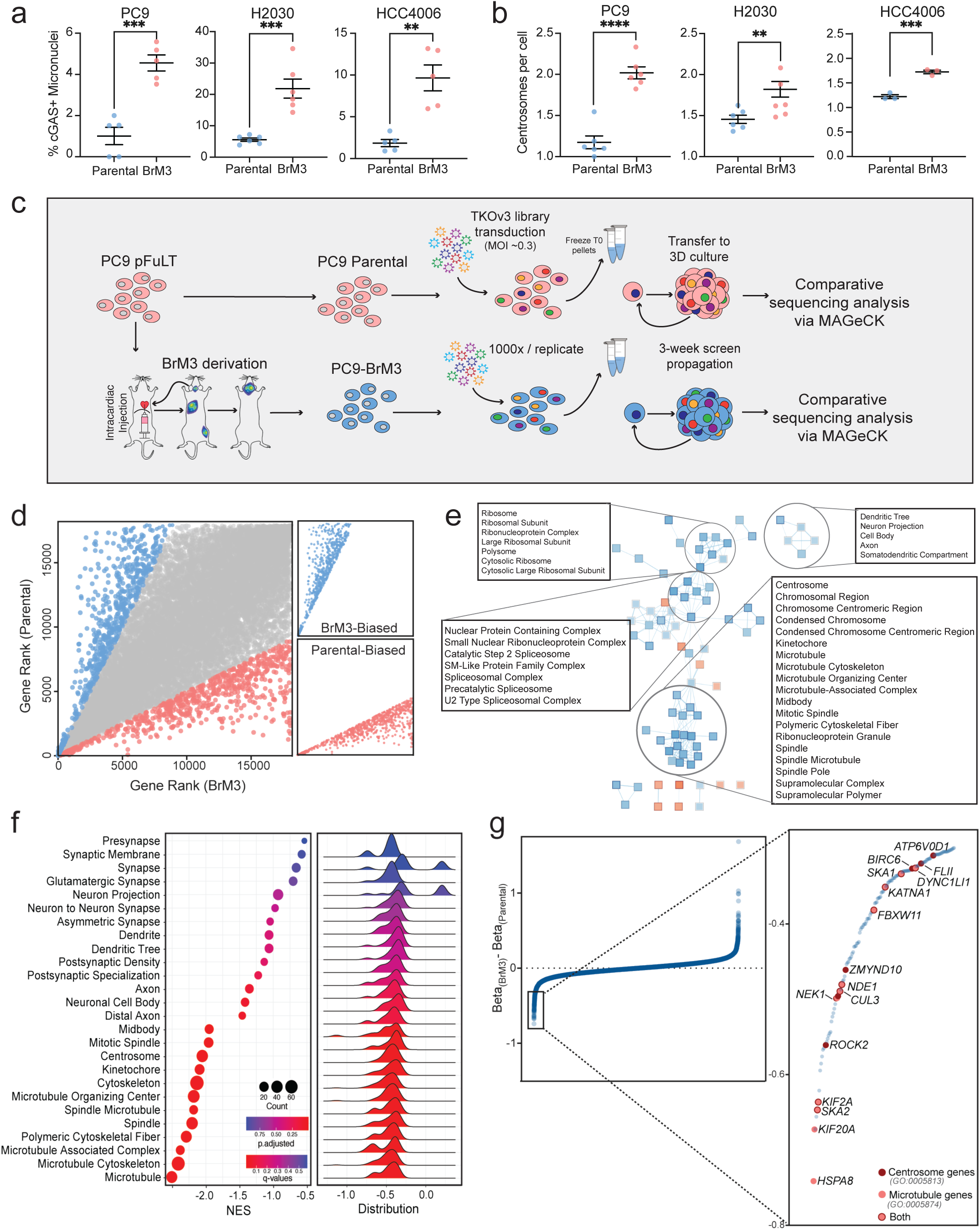
CIN-high metastatic cells are dependent on mitotic regulators. **(A)** Quantified immunofluorescence images of parental and brain-metastatic derivatives (BrM) measuring percentage of cells in each image that possess cGAS+ micronuclei (n=>50 cells/image, 5 images per cell line). **(B)** Quantified immunofluorescence (as in (A)) measuring the number of centrosomes per cell in non-dividing cells (n= >50 cells/image, 5 images per cell line). **(C)** Schematic of 3D spheroid screen. **(D)** Results of CRISPR/Cas9 screen, comparing parental and metastatic gene rank. **(E)** Cytoscape analysis showing pathway intersection of GSEA signatures revealed from the screen. **(F)**. GSEA dot plot of depleted gene signatures indicating enriched pathways in PC9-BrM cells compared to PC9 cells. **(G)** Snake plot of the beta score difference between PC9 parental and BrM cells, with microtubule and centrosome hits highlighted, demonstrating biased dependence on these features in BrM cells.

### *NDE1* is a selective dependency of brain-metastatic cells

The gene encoding one such mitotic regulator, NudE neurodevelopment gene 1 (*NDE1*), was a PC9-BrM3-biased dependency that emerged from the screen (Fig. 2a). *NDE1* encodes a direct regulator of cytoplasmic dynein that acts, at least in part, by recruiting the dynein adaptor protein LIS1. NDE1 has previously been implicated in mitosis and neuronal migration during development [22–25]. In contrast to other mitotic regulators previously implicated in cancer (Fig. 1g) [26–30], to our knowledge, *NDE1* has only been explored as a cancer biomarker, not as a cellular dependency [31]. Rather, *NDE1* has been extensively studied in the context of brain development, a context in which germline mutations result in neurological defects like lissencephaly and microcephaly. Homozygous *NDE1* mutant mice display neural migration defects, mitotic arrest, disordered microtubule organization, and increased levels of DNA damage, while cells lacking NDE1 display mitotic defects consistent with the phenotypes seen *in vivo* [22, 24, 25, 32–34]. Across lung cancer cell line models, brain-metastatic derivatives were strikingly more sensitive to *NDE1* knockout compared to parental cell lines in growth assays (Fig. 2b, S4a-c,e) and clonogenic analyses (Fig. 2c, S4d). To understand the relationship between *NDE1* expression and cancer progression in human patients, we queried *NDE1* expression levels in either normal, tumor, or metastatic tumor tissue across patient databases. Compared to adjacent normal tissue, human tumors, on average, expressed increased levels of *NDE1.* This expression was further increased in metastatic tumors analyzed by The Cancer Genome Atlas (TCGA) (Fig. 2d). *NDE1* knockout impaired the ability of PC9-BrM3 cells to colonize the brain *in vivo*, leading to increased overall survival, while not significantly affecting the growth of parental PC9 cells after intracardiac injection (Fig. 2e-f, S4f-g). Interestingly, of the mice that exhibited brain metastasis in the *NDE1*-knockout condition, two of the three mice that were profiled harbored tumors that had retained expression of NDE1, underscoring its importance in the survival of brain-metastatic cells (Fig. 2g-h). Together, these findings demonstrate that *NDE1* is a conserved, selective dependency in brain-metastatic models.

**Fig. 2.**
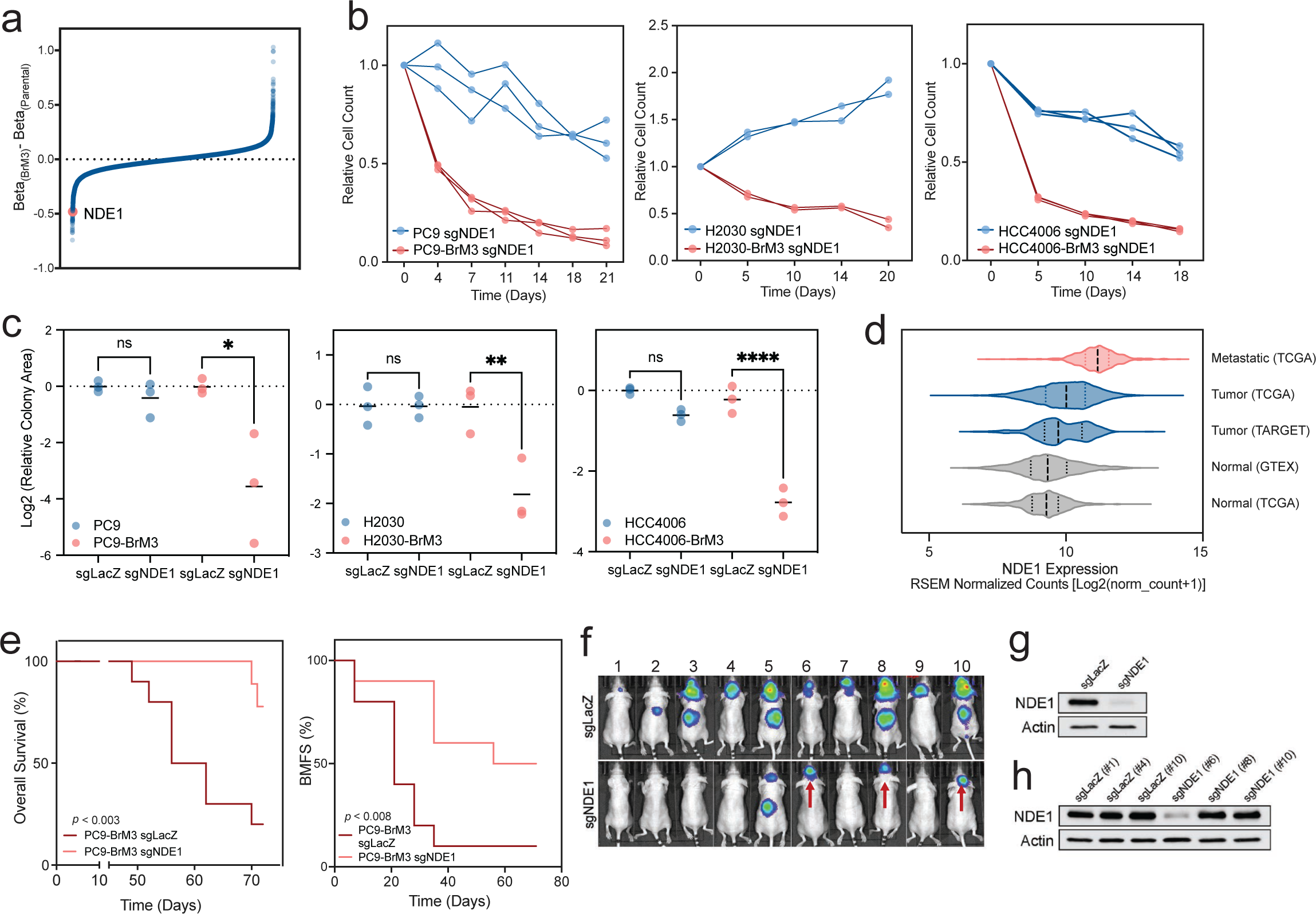
*NDE1* is a selective dependency of brain-metastatic cells. **(A)** Metastatic CRISPR screen data represented as a snake plot of the beta score difference between PC9 parental and BrM cells, with *NDE1* highlighted. **(B)** Longitudinal cell counting assays comparing cell growth between parental cells with *NDE1* knockout and BrM cells with *NDE1* knockout, normalized to LacZ targeting sgRNA controls. **(C)** Clonogenic analysis comparing colony-forming ability in parental cells to BrM cells with either LacZ targeting sgRNA control or *NDE1* knockout. **(D)** Normalized expression of *NDE1* in normal, tumor, and metastatic tumor tissues from selected cancer patient tissue databases. **(E)** Kaplan-Meier curves of overall survival in mice injected intracardially with PC9-BrM cells transduced with sgRNAs against LacZ (control) or *NDE1*. **(F)** Representative bioluminescent images of mice as quantified in (E). Red arrows indicate brain metastases that were collected at the end of the study and analyzed via western blot in Figure 2H. **(G-H)** Immunoblot analysis of NDE1 protein level in PC9-BrM3 cells at the population level, pre-intracardiac injection **(G)** and post-intracardiac injection, by mouse **(H)**.

### CIN is sufficient to confer *NDE1* dependence through increased mitotic dysregulation

Given the role of *NDE1* in mitotic regulation [22, 33], we hypothesized that dependence on *NDE1* was due to underlying features of chromosomal instability present in the brain-metastatic models. To determine if increased CIN is sufficient to drive *NDE1* dependence, we treated parental PC9 cells with pharmacologic agents previously known to induce chromosomal instability, including reversine, an Mps1 and Aurora kinase inhibitor, nocodazole, a microtubule polymerization inhibitor, and paclitaxel, a microtubule stabilizer. After treatment with a short pulse (<24 hours) and subsequent washout of each drug, cells displayed increased cGAS+ micronuclei, consistent with an induction of CIN. Regardless of the pharmacologic agent used, an increase in chromosomal instability was sufficient to sensitize parental PC9 cell lines to *NDE1* depletion (Fig. 3a-c). Of note, sensitivity to *NDE1* loss was proportional to the degree of CIN induction in each model, as measured by cGAS+ micronuclei. To understand how depletion of *NDE1* contributed to altered mitosis, we performed fixed and live cell imaging studies of PC9 and PC9-BrM3 cells with and without *NDE1* knockout. We observed a significant increase in phenotypes of mitotic dysregulation such as spindle multipolarity (Fig. 3d-f) and a drastic increase in time spent in mitosis (Fig. 3g-h), specifically in PC9-BrM3 cells upon *NDE1* knockout. While time in mitosis was generally extended in PC9-BrM3 cells lacking *NDE1*, there were a subset of cells that did not complete a successful division for greater than 500 minutes after DNA condensation. In contrast, parental PC9 cells divided on an average timescale of 50 minutes, even with *NDE1* depleted (Fig. 3g-j). In sum, these data are consistent with the hypothesis that increased chromosomal instability levels render brain-metastatic cells sensitive to *NDE1* loss through the buildup of mitotic defects associated with loss of mitotic regulation.

**Fig. 3.**
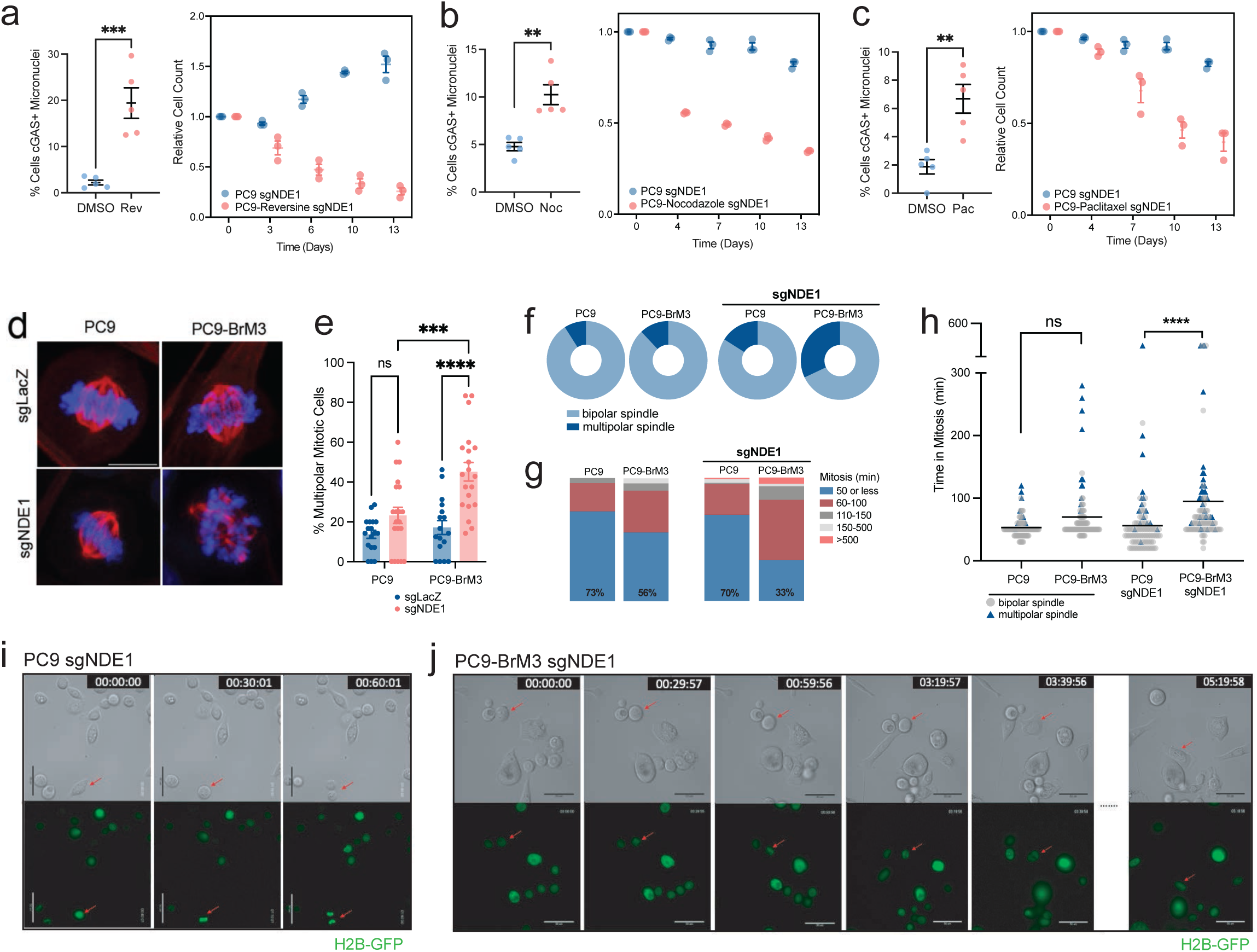
Increased CIN is sufficient to confer *NDE1* dependence. **(A-C)** Chromosomal instability was induced in parental PC9 cells using Reversine (A), Nocodazole (B), or Paclitaxel (C), and dependency on *NDE1* was measured using longitudinal cell counting. **(D**) Representative images showing multipolarity in BrM cells during mitosis, with exacerbated phenotypes in the *NDE1* knockout population quantified in **(E)**. **(F)** Pie chart showing the percentage of cells analyzed in (E) that underwent multipolar division. **(G-H)** Quantification of live cell imaging analysis, measuring the time a cell took to progress from DNA condensation to daughter cell separation. **(I)** Brightfield and fluorescent live-cell imaging time course of PC9-sgNDE1 cells, with DNA visualized using constitutive expression of a GFP-tagged histone 2B. The red arrow indicates the mitotic cell being followed. **(J)** Live cell imaging time course (as in (I)) with PC9-BrM3-sgNDE1 cells.

### STAG2 loss leads to increased chromosomal instability and sensitivity to *NDE1* depletion

To gain mechanistic insights into genetic and cell state differences between PC9-BrM3 cells and parental cells, we subsequently surveyed parental-biased genes that emerged from our whole genome screen. *STAG2*, *PAXIP1*, and *PAGR1* were striking genetic dependencies in PC9 cells but not in PC9-BrM3 lines (Fig. 4a). *STAG2*, *PAXIP1*, and *PAGR1* are members of an axis responsible for transcriptional regulation and tethering of sister chromatids during mitosis [35]. *STAG2* is frequently mutated in multiple cancer types, and loss of STAG2 function was also shown to increase metastatic potential in EWS-FLI-driven sarcoma models [36–38]. The three related proteins, which are strongly coessential in DepMap [39, 40], were recently characterized as tumor suppressive in lung cancer [41]. In validation cell viability assays, knockout of *STAG2*, *PAXIP1*, or *PAGR1* led to a drastic reduction in viability in PC9 cells, with little effect on viability of PC9-BrM3 cells (Fig. 4b). We hypothesized that the STA2/PAXIP1/PAGR1 axis may thus be less active in metastatic cells, rendering PC9-BrM3 cells insensitive to loss of each gene. Consistent with this, RNAseq data comparing PC9 cells to PC9-BrM3 cells showed a modest but significant reduction in expression of each gene (Fig. 4c). To determine whether genes related to STAG2 might be suppressed in the PC9-BrM3 cells, we created a signature of the top 100 gene expression correlates with *STAG2*, *PAXIP1*, and *PAGR1* across cell lines in the CCLE database [39, 40]. Using these respective expression signatures, we performed GSEA on our RNAseq data comparing PC9 and PC9-BrM3 cell lines. Gene signatures associated with *STAG2*, *PAXIP1*, and *PAGR1* were significantly downregulated in brain-metastatic cells compared to parental cells (Fig. 4d). STAG2 was also reduced at the protein level, specifically in BrM cells (Fig. 4e). In addition to regulating transcription, STAG2 is part of the cohesin complex, responsible for proper attachment and separation of sister chromatids during mitosis. Consistent with this role, loss of STAG2 has been previously shown to increase chromosomal instability and mitotic dysregulation in cancer cells [42, 43]. We sought to determine whether loss of STAG2, or the associated PAXIP1 and PAGR1, may be sufficient to increase CIN in parental PC9 cells to levels similar to those of the PC9-BrM3 derivative lines. In clonally derived *STAG2* and *PAXIP1* knockout PC9 cells, we observed visual phenotypes associated with mitotic dysregulation and increased CIN, including a significant increase in cGAS+ micronuclei and time spent in mitosis, compared to LacZ controls (Fig. 4f-h). Interestingly, knockout of *PAGR1* produced an insignificant increase in hallmark measures of CIN, indicating that the mechanism of PAGR1 sensitivity loss may be different. PC9 cells lacking *STAG2* or *PAXIP1* with a concomitant increase in CIN were sensitized to NDE1 depletion, similar to what was observed in brain-metastatic BrM cells (Fig. 4i). The relationship between the level of CIN induced by these alterations, measured via cGAS+ micronuclei, and sensitivity to *NDE1* loss was profoundly correlated in the PC9 knockout clones, highlighting the relationship between chromosomal instability and *NDE1* genetic dependence (Fig. 4j). Together, these data suggest that disruption of the STAG2/PAXIP1/PAGR1 axis through decreased activity or expression may drive the increased CIN observed in PC9-BrM3 cells.

**Fig. 4.**
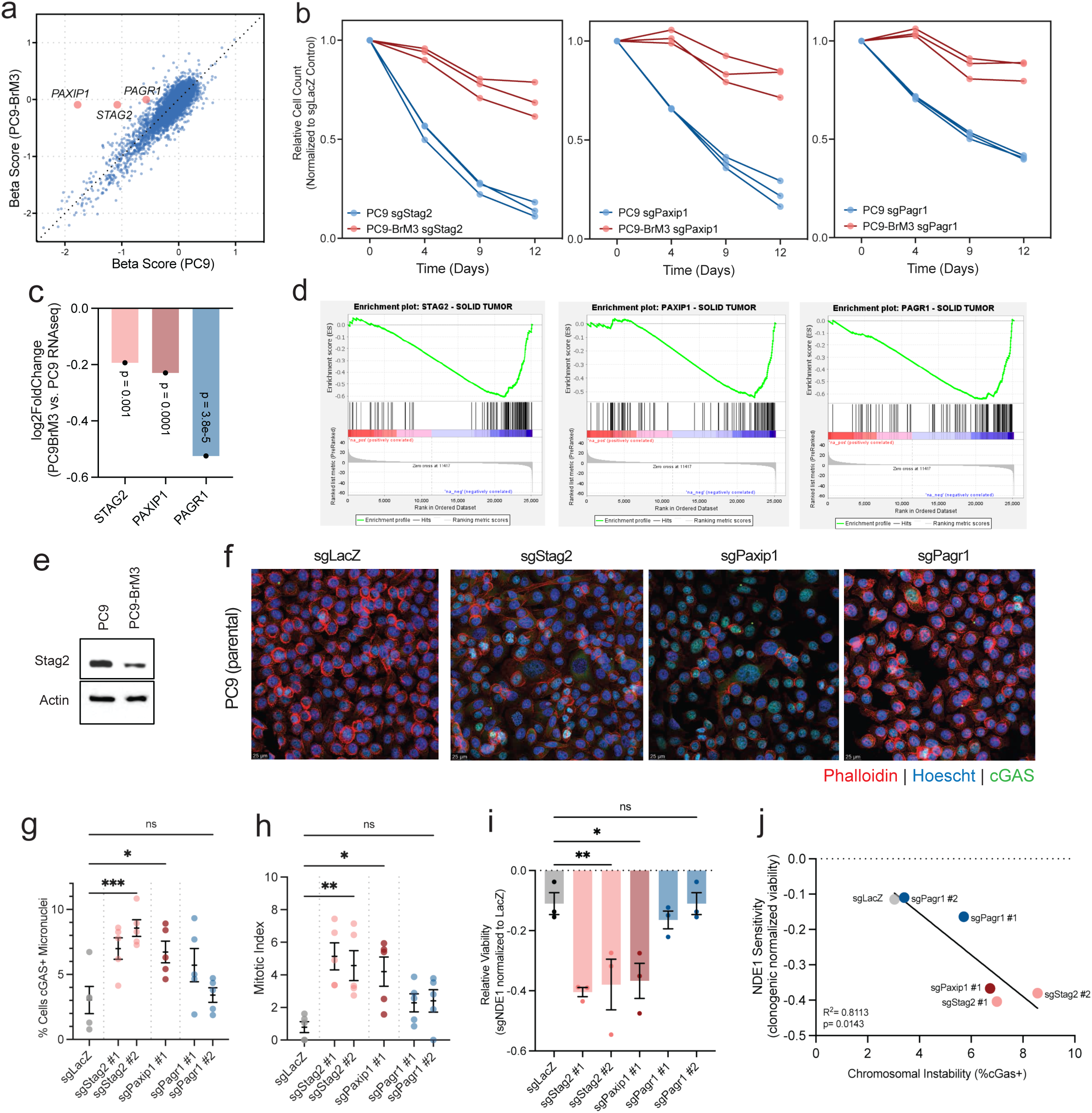
Loss of *STAG2* leads to increased CIN and sensitivity to *NDE1* knockout. **(A)** Metastatic CRISPR screen data represented as a comparison of beta-scores between parental and BrM cells, with *STAG2*, *PAXIP1*, and *PAGR1* highlighted. **(B)** Longitudinal cell counting assays comparing cell growth of parental PC9 cells with PC9-BrM3 cells after knockout of *STAG2*, *PAXIP1*, and *PAGR1*, normalized to LacZ targeting sgRNA controls. **(C)** Log2 fold change of *STAG2*, *PAXIP1*, and *PAGR1* expression from RNA-seq comparing PC9-BrM3 to PC9 parental cells. **(D)** GSEAPrereanked results showing depletion of *STAG2*, *PAXIP1*, and *PAGR1* expression signatures in RNAseq data comparing differentially expressed genes in PC9 and PC9-BrM cells. **(E)** Western blot comparing STAG2 expression between PC9 and PC9-BrM3 cells. **(F)** Representative immunofluorescence images staining for cGAS+ micronuclei in PC9 parental cells with *STAG2*, *PAXIP1*, and *PAGR1* knockout, with **(G)** depicting quantification of data from multiple images (n= >50 cells/image, 5 images per cell line). **(H)** Quantification of mitotic index (% of cells actively undergoing mitosis) in images like those shown in (F) (n= >50 cells/image, 5 images per cell line). **(I)** Cell counting assay measuring relative viability of *STAG2*, *PAXIP1*, and *PAGR1* knockout clones after *NDE1* knockout, normalized to LacZ. **(J)** Linear regression of NDE1 sensitivity and CIN, as measured by cGAS+ micronuclei.

### Chr17p loss is independently sufficient to confer *NDE1* dependence

To determine if there are conserved features of *NDE1*-dependent cell lines beyond those found in the context of our screen, we queried the DepMap [39, 40] database for possible correlations with *NDE1* gene dependence. Strikingly, *NDE1* dependence is highly correlated with the loss of chromosome-arm 17p across cancer types (Fig. 5a). When we queried the degree of correlation between *NDE1* dependence and the copy number status of all genes in the genome, the top 88 associations were copy number losses in genes that exclusively map to chromosome Chr17p (Fig. 5b). Reciprocally, across the genome, *NDE1* is the genetic dependency most highly correlated with Chr17p loss (Fig. 5c). This finding was of particular interest because Chr17p is commonly lost in human cancers as a way to inactivate *TP53*, and a recent study has shown an enrichment of Chr17p loss in brain metastases [12]. We validated this finding experimentally across a selection of cell lines annotated as harboring Chr17p loss, copy number neutrality, or gain in DepMap [39, 40], and observed heightened sensitivity to *NDE1* depletion in cell lines containing Chr17p loss (Fig. 5d). Furthermore, partial loss of chromosome Chr17p was sufficient to confer *NDE1* dependence in a previously described isogenic model in which Chr17p loss was induced by CRISPR/Cas9 in DU145 prostate cancer cells (DU145-Chr17p-del) (Fig. 5e) [44]. Interestingly, wild-type and DU145-Chr17p-del cells had similar levels of cGAS+ micronuclei, suggesting that Chr17p loss is independently sufficient to confer *NDE1* dependence, even if CIN levels are not elevated (Fig. 5f). To determine if the presence of both increased CIN and Chr17p loss confers additive dependence on *NDE1*, we treated WT and DU145-Chr17p-del cells with Reversine to increase CIN as previously described (Fig. 3). While Chr17p loss and CIN induction were each sufficient to increase *NDE1* dependence on their own, cells with both Chr17p loss and Reversine treatment were further sensitized to *NDE1* depletion (Fig. 5g). Interestingly, in lung cancer patients included in the MSK-Met sequencing cohort [5], Chr17p loss was associated with an increased probability of developing brain metastasis (Fig. 5h). In this cohort, the presence of any arm-level chromosome alteration biased patients towards brain metastasis, with Chr17p loss being within the most enriched arm losses found in CNS metastases (Fig. 5i). Thus, *NDE1* dependence is tightly linked to Chr17p loss, potentially independent of CIN.

**Fig. 5.**
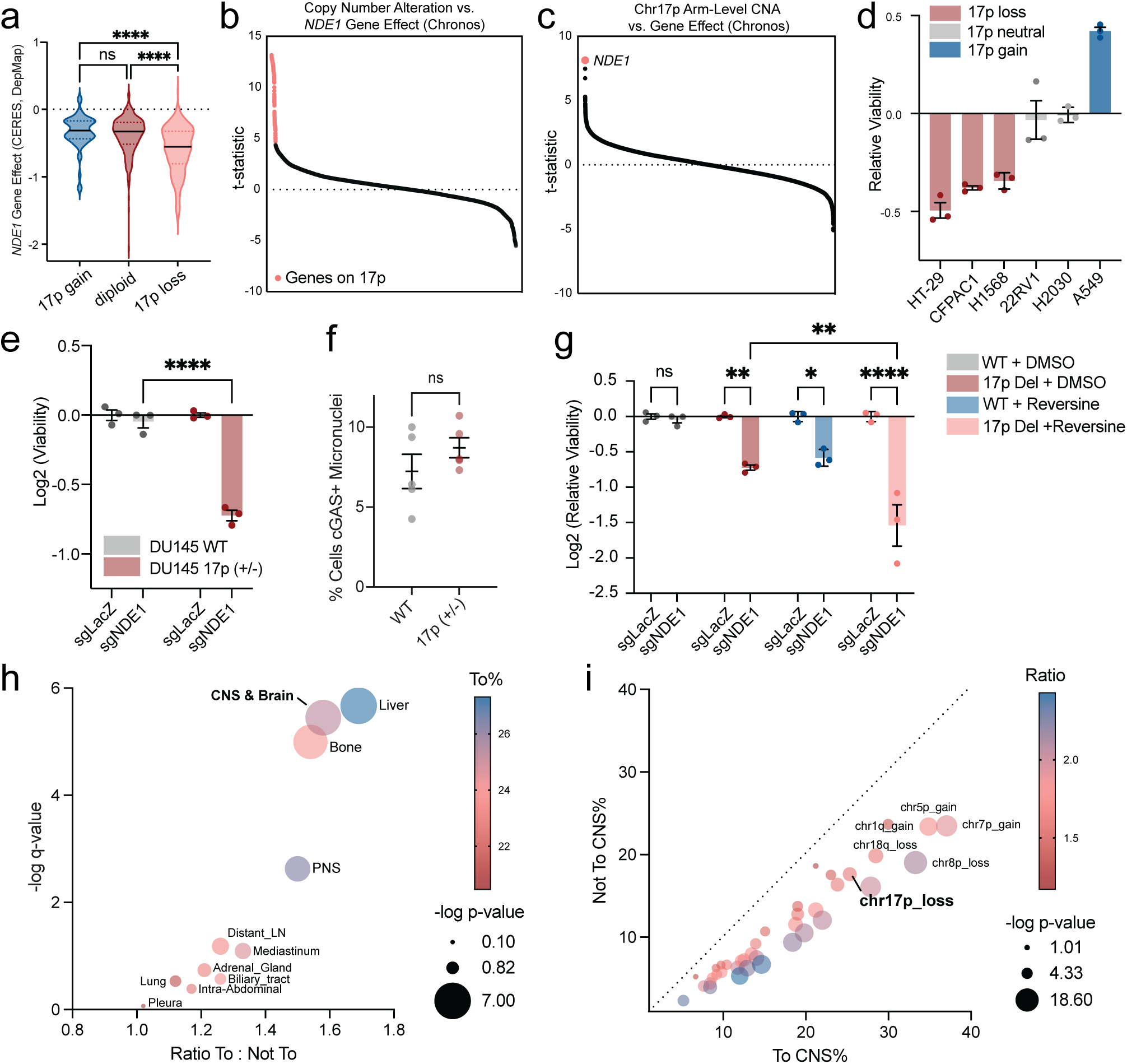
Chr17p loss is independently sufficient to confer *NDE1* dependence. **(A).** Violin plot comparing *NDE1* gene effect across cancer cell lines annotated as either Chr17p gain, Chr17p neutral, or Chr17p loss in DepMap. **(B)** Correlation of copy number across genes in the genome to *NDE1* gene knockout effect across cell lines in DepMap, represented as a snake plot ranked by t-statistic. Genes on Chr17p are annotated in pink. **(C)** Correlation of gene knockout effects across the genome to Chr17p arm-level copy number across cell lines in DepMap, represented as a snake plot ranked by t-statistic. *NDE1* is highlighted. **(D)** Relative viability of selected cell lines with annotated Chr17p status after *NDE1* knockout using CRISPR/Cas9, normalized to LacZ targeting sgRNA controls. **(E)** Cell counting assay comparing cell viability after knockout of *NDE1* in DU145 WT vs. DU145-Chr17pdel cell lines. **(F)** Quantified immunofluorescence images of DU145 WT vs. DU145-Chr17pdel cell lines measuring percentage of cells in a given image that possess cGAS+ micronuclei (n=>50 cells/image, 5 images per cell line). **(G)** Cell counting assay comparing cell viability after *NDEI* knockout in DU145 WT vs. DU145-Chr17pdel cell lines, with or without Reversine pre-treatment to induce CIN. **(H)** For lung cancer patients in the MSK-met database, the ratio of metastasis to a specific organ site when the patient had annotated Chr17p loss, versus no metastasis to that organ site when the patient had Chr17p loss. Higher ratios indicate more metastasis to a specific organ site among patients with Chr17p loss. **(I)** Bias plot comparing the percentage of lung cancer patients with a given arm-level alteration that metastasized to the CNS. The dotted line represents where points would fall if an equal percentage of patients were positive and negative for CNS metastasis when possessing the given arm loss alteration.

### Chr17p loss confers *NDE1* dependence through partial loss of *NDEL1*

Because heterozygous loss of Chr17p is a common way to inactivate p53, we initially hypothesized that the *NDE1* dependence observed in Chr17p-deleted cancer cells was caused by p53 inactivation. However, multiple lines of evidence pointed away from this conclusion: (1) PC9 parental cells and their brain-metastatic derivatives have inactivating mutations in *TP53* on both alleles (R248Q), making it unlikely that differences resolved in the screen were due to differential p53 activity; (2) each of our cell line models and their matched brain-metastatic derivatives were equally resistant to reactivation of p53 with the MDM2 inhibitor Nutlin (Fig. S5a); and (3) depletion of *TP53* was insufficient to induce *NDE1* dependence in both parental PC9 cells and h-TERT transformed retinal pigment epithelial cells (RPE-hTERT), which have intact p53 (Fig. S5b). Additionally, *TP53* mutation status did not correlate with *NDE1* dependence in DepMap (Fig. S5c). Given these results, we reasoned that loss of other genes on Chr17p were likely responsible for the increased *NDE1* dependence observed in Chr17 loss models. Interestingly, a paralog of *NDE1*, NudE, Neurodevelopment-Protein 1 Like 1 (*NDEL1*), is located on chromosome 17p (Fig. 6a), and its loss has been shown to exacerbate the effects of *NDE1* loss [23]. Indeed, *NDE1* dependence was correlated with expression and copy number of *NDEL1* across cell lines in DepMap (Fig. 6b-c) [39, 40]. To test the hypothesis that partial loss of *NDEL1* is sufficient to drive *NDE1* dependence, we derived clones of PC9 cells with partial knockout of *NDEL1* using CRISPR/Cas9 to model heterozygous loss (Fig. 6d). Both clones with partial loss of *NDEL1* (PC9-sgNDEL1-C8 and C9) were dramatically sensitized to *NDE1* deletion (Fig. 6e). Further, approximately 50% knockdown of NDEL1 levels using a dox-inducible shRNA was also sufficient to sensitize PC9 cells to *NDE1* deletion (Fig. S5d-e). In cell lines with annotated Chr17p loss (from Fig. 5d), re-expression of NDEL1 largely rescued *NDE1*-dependence (Fig. 6f-g). A similar rescue was observed in the DU145-Chr17p-del model (from Fig. 5e), which displayed about 50% NDEL1 expression at baseline compared to WT (Fig. 6h-i). Notably, another gene of interest, *PAFAH1B1* (LIS1), which encodes a binding partner of NDE1 and is also located on chromosome Chr17p, was insufficient to drive *NDE1* dependence when its expression was reduced with shRNA (Fig. S5f). Further, in contrast to *NDEL1*, re-expression of Lis1 did not rescue sensitivity to *NDE1* in cell lines with Chr17p loss (Fig. S5g). To understand the effects of paralog depletion *in vivo*, we infected PC9-sgNDEL1_C8 cells with a dox-inducible CRISPR/Cas9 system targeting *NDE1* and injected the cells into the flanks of athymic nude mice. Once tumors were established (100mm^3^) and *NDE1* knockout was induced, mice containing tumors with partial *NDEL1* deletion exhibited significantly decreased tumor growth over the course of the study (Fig. 6j-k). Taken together, these data demonstrate that partial loss of *NDEL1* drives induced *NDE1* dependence in the context of Chr17p loss.

**Fig. 6.**
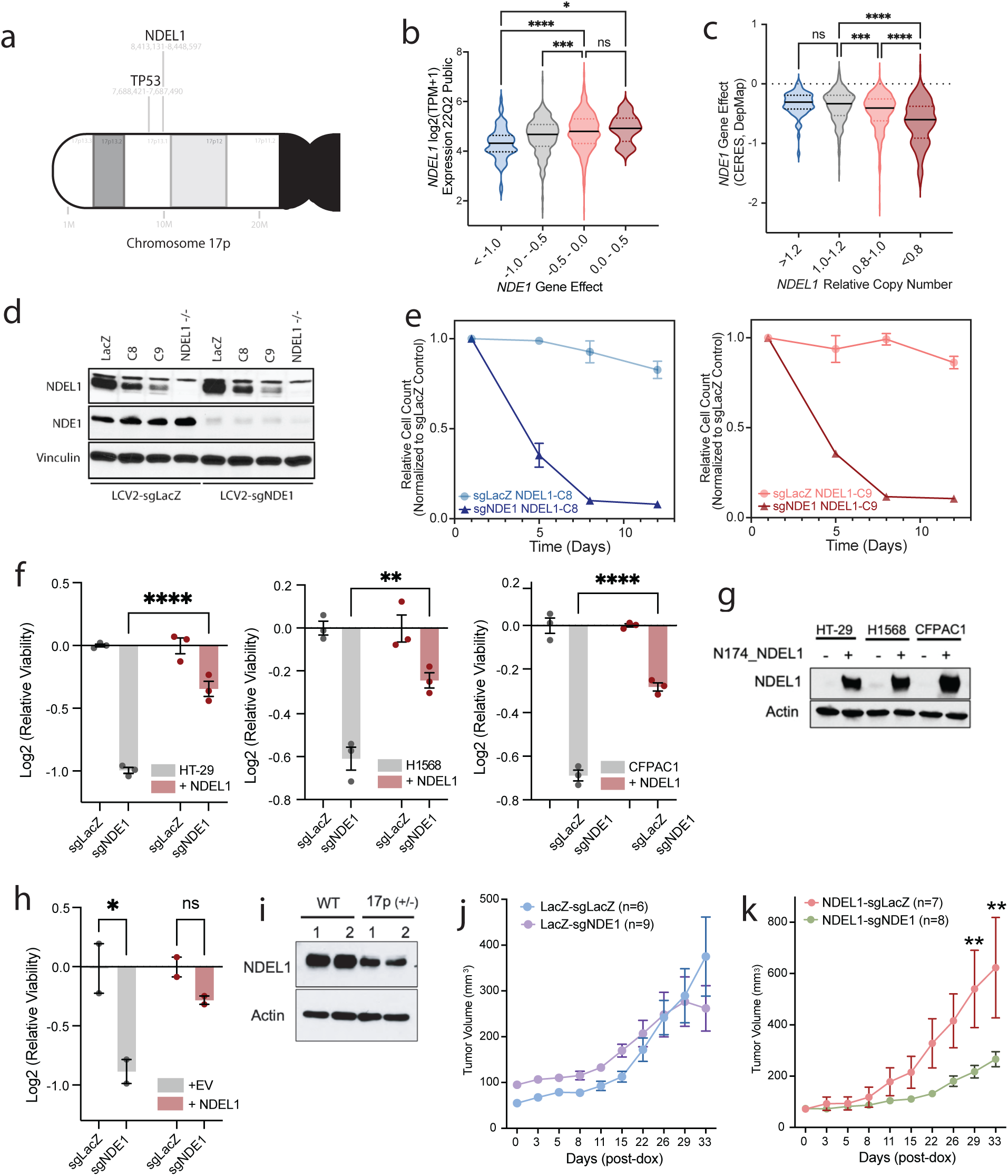
Chr17p loss confers *NDE1* dependence through partial loss of *NDEL1*. **(A)** Schematic of *TP53* and *NDEL1* gene location on chromosome Chr17p. **(B)** Violin plot comparing *NDEL1* expression to *NDE1* gene effect across cancer cell lines in DepMap. **(C)** Violin plot comparing *NDEL1* relative copy number to *NDE1* gene effect across cancer cell lines in DepMap. **(D)** Western blot analysis of *NDEL1* partial knockout clones (PC9-C8 and PC9-C9) compared to a complete *NDE1* knockout, all generated with CRISPR/Cas9. **(E)** Longitudinal cell counting assays measuring cell viability in *NDEL1* partial knockout clones after *NDE1* knockout, normalized internally to LacZ targeting sgRNA controls. **(F)** Rescue of *NDE1* sensitivity after overexpression of *NDEL1* in cell lines with annotated Chr17p loss in DepMap, with overexpression confirmed in **(G)**. **(H)** Same as (F), in the isogenic DU145-Chr17pdel model. **(I)** Western blot analysis showing partial NDEL1 expression in the DU145-Chr17pdel cell line compared to DU145-WT. 1 and 2 are independent cell harvests of asynchronous cell populations, separated by 14 days. **(J)** Tumor volumes from nude mice injected with PC9 parental cells, after dox-inducible knockout of *NDE1* or LacZ control using CRISPR/Cas9. **(K)** same as (J) with PC9 NDEL1-C8 cells injected.

In sum, our findings suggest a model by which increased CIN (induced by mechanisms that can include loss of STAG2) and Chr17p loss (through the partial deletion of *NDEL1*) independently drive *NDE1* dependence in the context of metastasis. Interestingly, these phenomena often co-occur, as Chr17p loss, though inactivation of p53, can increase cellular tolerance of CIN, while CIN can facilitate the generation and subsequent selection for Chr17p loss [10, 11, 45]. When combined, these reinforcing features endow an additive dependency on *NDE1* in metastatic cancer cells (Fig. 7).

**Fig. 7.**
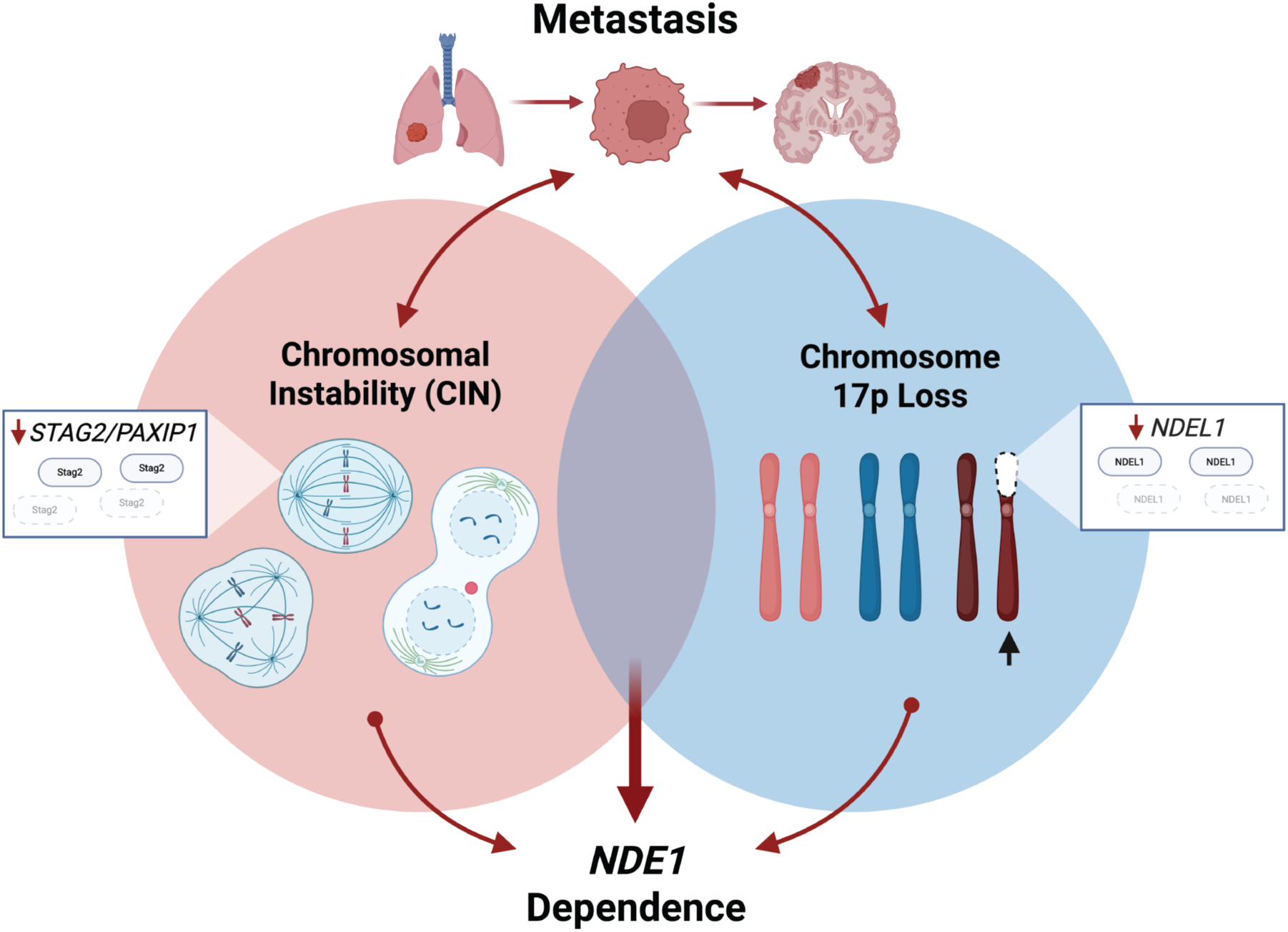
Schematic: Chromosomal instability and chromosome 17p loss drive convergent *NDE1* synthetic lethality in metastatic cancer cells.

## Discussion

In this study, we uncover a set of collateral sensitivities resulting from metastasis to distal organ sites such as the brain. One such collateral sensitivity, to loss of *NDE1*, is independently driven by two recurrent features of metastatic cells: high levels of CIN and partial deletion of its paralog gene, *NDEL1*, in the context of heterozygous Chr17p loss. Interestingly, we show that the presence of both features, which commonly co-occur in metastatic cancers, including lung cancer brain metastases [5, 6, 12], results in compounded *NDE1* sensitivity. Therefore, we implicate NDE1 as a novel ‘dual’ synthetic lethal vulnerability imposed by two separate but interrelated features important for the evolution of metastasis. This concept of a dual synthetic lethality extends previous studies implicating partial chromosome losses as underleveraged drivers of tumor progression, which often result in collateral sensitivities [44, 46].

While many studies have previously shown that chromosomal instability is associated with metastasis and progression in patients, few have characterized tumor cell intrinsic vulnerabilities associated with a CIN-high state. Our work builds upon previous studies which have implicated the spindle assembly checkpoint (SAC) members (*BUB1B* and *MAD2L1*) and *KIF18A* as core dependencies associated with whole-genome doubling, aneuploidy, and the maintenance of CIN [21, 47, 48]. These studies suggest a similar model in which depletion of mitotic regulators causes deleterious mitotic defects, specifically in CIN-high cells with abnormal DNA content, the accumulation of which drives the loss of cell viability. In each case, knockout of the dependency gene largely spares cells that are CIN-low or diploid. Here, we demonstrate that NDE1, an understudied regulator of dynein function and microtubule organization involved in proper chromosome segregation, is an additional selective vulnerability present in cancer cells with underlying chromosomal instability, particularly in the context of brain metastasis. We demonstrate that, similar to *KIF18A* and genes encoding members of the SAC, depletion of *NDE1* leads to loss of mitotic regulation, an increase in the percentage of cells undergoing multipolar division, and drastic extension of mitoses.

While the relationship between *NDE1* dependence and CIN is largely consistent with previous models, the connection between *NDE1* dependence and Chr17p loss has, to our knowledge, not been previously explored despite its strong correlation in DepMap [39, 40] and the prevalence of Chr17p loss across cancer types [10]. Though our metastatic-derivative models did not show increased selection for Chr17p loss, patient data from MSK-Met, as well as additional studies, indicate that Chr17p loss is highly enriched specifically in brain metastasis, likely because heterozygous loss of Chr17p is a common mechanism for inactivation of p53 after a single mutation has been acquired in the other copy. Despite the increased fitness associated with p53 inactivation, we show that cells with heterozygous loss of Chr17p acquire dependency on *NDE1* due to decreased expression of its paralog, *NDEL1*, which is chromosomally adjacent to *TP53*. Because loss of Chr17p is also often a ‘gateway’ to chromosomal instability through p53 inactivation, we propose that targeting NDE1 may be especially effective in cancers with this alteration.

Additional results from our screen suggest that downregulation of the STAG2/PAXIP1/PAGR1 axis may contribute to the acquisition of CIN during tumor metastasis. Recent studies showing that epithelial-to-mesenchymal (EMT) transition causes durable CIN in pancreatic cancer highlight the importance of CIN in progressive malignancy [49]. With the relationship between EMT and CIN now firmly established, it has not escaped us that a larger cell state, such as EMT, may be responsible for the increased CIN and depletion of the STAG2/PAXIP1/PAGR1 axis found in our brain-metastatic derivative cell lines, though this was not directly explored. Previous studies have also implicated loss of the cohesion complex as a key feature of widespread transcriptional change, as is necessary during EMT, though this is less established [38, 41, 50]. Elegant studies in other cancer types, such as EWS-FLI-driven sarcoma, demonstrated that loss of *STAG2* is sufficient to increase metastatic potential in a tail-vein injection model [38]. Additionally, loss of *STAG2*, as is common in bladder cancer, is associated with increased metastasis and worse clinical outcomes [37]. These findings, combined with the results of our whole-genome screen, suggest that suppression of the STAG2/PAXIP1/PAGR1 axis is one biologically viable way to acquire traits important for metastatic evolution, such as increased CIN.

Important limitations of this study center around the challenges of large-scale genomic screening in the context of cancer metastasis. While our parental and metastatic-derivative cell line models recapitulate key features of metastasis seen in patients, like increased CIN, the endpoint-to-endpoint nature of the isogenic system prevents us from determining the stage of the metastatic cascade that most contributes to acquired *NDE1* dependence. However, it is clear that dependency on *NDE1* remains durable upon subsequent passage of the cell lines *in vitro* and is likely not directly dependent on the microenvironment of the brain. Though not directly tested in this study, we hypothesize that dependence on *NDE1* may extend to other sites of metastatic tropism, provided CIN is increased and/or chromosome Chr17p is lost in these cells.

In totality, our study reveals that the process of metastatic evolution can confer cell-autonomous dependencies on cancer cells. Further, it identifies the first example, to our knowledge, of a convergent synthetic lethal vulnerability driven by two traits of a cancer cell. These studies suggest that in the future, it may be possible to design therapeutic strategies that specifically target the metastatic lesions that drive cancer mortality.

## Online Methods

### Study Approval

The Institutional Animal Care and Use Committee at Duke University (IACUC) approved all animal procedures and studies. The study’s IACUC protocol number is A189-22-11.

### Chemicals

#### In Vitro

Doxycycline was purchased from VWR (103516-794) and was prepared at 2 mg/mL in molecular biology-grade water. Puromycin Dihydrochloride (6136) was purchased from Gold Biotechnology. Blastocidin S was purchased from Sigma (203351).

#### In Vivo

Doxycycline hyclate 98% (AC446061000) was purchased from Thermo Fisher Scientific.

### Plasmids

The following plasmids were purchased from Addgene: lentiCRISPR v2 (LCV2, #52961), Tet-pLKO-puro (#21915), pRDA_355 (#187159), pKLV2-EF1a-Cas9Bsd-W (#68343), N174-MCS (#81061), psPAX2 (#12260), pMD2.g (#12259), pSpCas9(BB)-2A-GFP (PX458, #48138), pcDNA3 Ndel1 (#12571) pcDNA3 Lis1 (#12574).

### Cell Culture

The following cell lines were used: PC9 (RRID: CVCL_B260), H2030 (RRID: CVCL_1517), HCC4006 (RRID: CVCL_1269), HEK-293-FT (RRID: CVCL_6911), HT-29 (RRID: CVCL_0320), CFPAC-1 (RRID: CVCL_1119), NC1-H1568 (RRID: CVCL_1476), 22Rv1 (RRID: CVCL_1045), A549 (RRID: CVCL_0023). RPE-hTERT (RRID: CVCL_4388), and DU145 (RRID: CVCL_0105). RPE-hTERT cells were a gift from Dr. Beth Sullivan (Duke University). DU145 cells and DU145 17pDel cells were a gift from Xinna Zhang and Xiongbin Lu (Indiana University). All other cell lines were purchased from the Duke University Cell Culture Facility or ATCC and were grown in a humidified incubator at 37°C with 5% CO_2_. Before use, cell line authenticity was confirmed through short tandem repeat profiling performed at the Duke University DNA Analysis Facility. Cells were maintained in culture for a maximum of 6 weeks before being discarded. Testing with the MycoAlert Mycoplasma Detection Kit (Lonza, # LT07-318) was performed regularly to ensure the cell lines were free from *Mycoplasma* contamination. All cell lines were cultured in RPMI (Gibco), except RPE-hTERT cells, which were cultured in DMEM:F12 (Gibco), and 293FT cells, which were cultured in DMEM high glucose (Gibco) with 1% nonessential amino acids and 1% sodium pyruvate added. All media was supplemented with 10% fetal bovine serum (VWR) and 1% penicillin/streptomycin (Gibco).

### Intracardiac Injections

As previously described [52], 4×10^5^ lung cancer cells stably transduced with pFU-luciferase-Tomato (pFuLT) DNA were suspended in 100 uL PBS and injected into the left cardiac ventricle with a 30-gauge needle. Mice were anesthetized with 5% isoflurane prior to injections. Mice were monitored to ensure recovery from anesthesia and were imaged weekly to monitor for progression of disease burden using an IVIS XR bioluminescent imager. The presence of brain metastases was confirmed through *in vivo* BLI followed by isolation of brain metastases and cell culture expansion *in vitro*. Living Image software was used for analysis of BLI data. Age-matched, 8–12-week-old female athymic nu/nu mice were used for all IC injection studies (Jackson Laboratory). For *in vivo* metastasis experiments evaluating the phenotypic effects of CRISPR-guided gene knockouts, intracardiac injections of PC9-BrM3 cells lentivirally transduced with LCV2-LacZ were compared to intracardially-injected PC9-BrM3 cells transduced with LCV2-NDE1, KIF2A, or NEK1. Similarly, parental PC9 cells transduced with either LCV2-LacZ or LCV2-sgNDE1 were injected intracardially and measured via serial IVIS bioluminescent imaging to monitor disease progression.

### Derivation of Brain-Metastatic Lung Cancer Cell Lines

Generation of brain-tropic metastatic cell lines was performed as described previously [52, 53]. Human parental lung cancer cell lines PC9, HCC4006, and H2030 expressing pFuLT were injected intracardially as described above under *Intracardiac Injections* and monitored weekly via IVIS bioluminescent imaging. Upon presentation with advanced brain metastatic burden (∼30 days post-intracardiac injection), animals were euthanized and whole brains were collected, minced, and digested in RPMI culture medium (Gibco) supplemented with 0.125% collagenase III (Thermo Fisher) and 0.1% hyaluronidase (Thermo Fisher) and placed on a rotator for 5 hours at room temperature. After centrifugation, cell pellets were further digested in 0.25% trypsin (Gibco) for 10 minutes at 37°C, then pelleted. Cells were then resuspended in culture media containing 1X Anti-Anti (Thermo Fisher) and expanded to near-confluence in a 10cm culture plate. Cells were then lifted with 0.25% trypsin and sorted for Tomato-positivity.

### RNAseq

Differential gene expression analysis of 3D-cultured PC9 parental and PC9-BrM3 cells was performed by extracting total cellular RNA from equally confluent cell cultures using the RNeasy RNA extraction kit (Qiagen). Libraries were sequenced by paired-end sequencing on an Illumina sequencer run by the Duke University Genome Sequencing facility. Raw RNA-seq fastq data files were processed using TrimGalore software followed by read quality assessment with the Fast-QC analysis tool. Trimmed and quality-filtered reads were then mapped to the GRCh37 version of the human genome and transcriptome using the HISAT2 aligner and kept for subsequent analysis if they mapped to a single genomic location. Gene counts were compiled using FeatureCounts software. Normalization and differential expression were carried out by DESeq2 in an R programming environment.

### Low-Pass Whole Genome Sequencing and Analysis

Total genomic DNA was extracted from cell pellets using the QuickDNA Miniprep Plus Kit (Zymo). For each cell line, two separate cell pellets were sequenced as biological replicates. Whole-genome sequencing was performed at the Duke Sequencing and Genomic Technologies Core on the Illumina NextSeq 1000 at 1X coverage to capture large-scale genomic alterations. Sequencing reads were aligned to the hg38 reference genome using bowtie2 (version 2.4.2) and default parameters. After alignment, arm-level alterations were identified using AneuFinder [3] (version 1.9.0) to generate genome-wide copy-number profiles. Reads were binned at 1 Mb resolution and segmented using the edivisive-algorithm.

### CRISPR/Cas9 Screen

PC9 or PC9-BrM3 cells were transduced via spinfection with Toronto CRISPR Human Knockout Library (TKOv3) lentivirus according to methods previously described [54]. Briefly, 3.84E8 cells of each cell line were spinfected with TKOv3 lentivirus in 6-well plates at a seeding density of 4E6 cells/well for 1 hour at 2250 rpm at 25°C. Following spinfection, cells were immediately transferred to 15 cm dishes at 8E6 cells/dish and incubated overnight at 37°C. 24 hours following library transduction, culture media was replaced with fresh media containing 2 ug/mL puromycin for selection. After 72 hours of puromycin exposure, cell counts were taken to confirm >1000x library coverage of the transduced population at an MOI of ∼0.25. Transduced cells were expanded in puromycin for a total of 7 days following spinfection, at which point the transduced cell population was collected and seeded into 3D cell culture conditions on ultra-low attachment plates (Corning). Cells were counted and replated every 3-4 days, for a total of 21 days. Throughout the duration of the screen, each replicate was represented by a minimum of 75E6 cells, sufficient to provide 1000x coverage of the library (∼1000 cells per unique sgRNA). Samples of 68-75E6 cells were collected when the screen was initiated (T_initial_), terminated (T_final_), and at each passage. Following screen completion, DNA was extracted with the QIAamp Blood Maxi Kit (Qiagen) according to the manufacturer’s specifications and prepared for sequencing as previously described [55].

### CRISPR/Cas9 Screen Analysis

Deep sequencing was performed on an Illumina Nextseq platform (75 bp, single-ended) to identify differences in library composition after guide enrichment or depletion from T_initial_ to T_final_. All sequencing was performed by the Duke Sequencing and Genomics Core. Raw sequencing reads were trimmed and processed, and analysis of enrichment and depletion metrics comparing PC9 parental to PC9-BrM3 cells was performed using the MAGeCK analysis platform under default settings [56]. A differential gene dependency score between parental PC9 and PC9-BrM3 cells from the CRISPR/Cas9 screen was determined for each gene by calculating the difference in beta scores between parental and metastatic cell conditions. Ranked gene lists sorted by this differential score were imported for gene set enrichment analysis (GSEA) (The Broad Institute [57]) using ClusterProfiler software. The pre-ranked gene list was processed under default settings and size filters for analysis across all signatures contained within the following mSigDB databases: Hallmark (v6.2, 50 gene sets), Positional (v6.2, 259 gene sets), Curated (v6.2, 3,648 gene sets), Motif (v6.2, 776 gene sets), Computational (v6.2, 782 gene sets), Gene Ontology (v6.2, 4,364 gene sets), Oncogenic Signatures (v6.2, 187 gene sets), and Immunologic Signatures (v6.2, 4,872 gene sets). Additionally, the pre-ranked gene list was imported into Cytoscape software (National Resource for Nework Biology [58]) for cell pathway network analysis under default settings.

### 3D Cell Culture

For the CRISPR screen and in select validation assays (Fig. 2b, S3d,g), cells were grown as tumor spheroids in 3D culture media containing RPMI, 10g/L methylcellulose powder (Fisher M-352), 10% fetal bovine serum (VWR) and 1% penicillin/streptomycin (Gibco). 3D media was prepared by first autoclaving dry methylcellulose powder in an empty cell culture container to ensure sterility prior to addition of 37°C pre-warmed culture media. The methylcellulose-media mixture was then immediately transferred to a 4°C coldroom and stirred on a stir plate continuously overnight to ensure that all methylcellulose was fully dissolved prior to use in cell culture. Cells were plated in ultra-low attachment culture dishes (Corning) to encourage non-adherent growth. To passage cells, a 1:1 volume of PBS was added to reduce viscosity of media before centrifugation. Tumor spheroids were then dissociated with Accutase (Stem Cell Technologies) at room temperature for 30 minutes and subsequently counted and replated at desired seeding densities.

### CRISPR/Cas9 Knockouts

#### LentiCRISPRv2 (LCV2)

sgRNAs against the chosen gene were cloned into LCV2 (Addgene #52961) and validated with Sanger sequencing. To generate lentivirus, LCV2 constructs were co-transfected with psPAX2 (Addgene #12260) and pMD2.g (Addgene #12259) packaging plasmids into AAVpro 293Ts (Takara) using Lipofectamine 2000 (Thermo) according to the manufacturer’s protocol. Following a 4-hour incubation, the transfection medium was aspirated and replaced with viral harvest medium containing 30% FBS. After 48 h or 72 h of viral generation, the virus-containing medium was collected, filtered with a 0.45 μm filter and stored at −80 °C. Cells were subjected to spinfection with a transduction mixture of virus, 8 μg/mL polybrene, and media at 2,250 rpm for 1 hour. The following day, the media was refreshed, and the cells with plasmid integration were selected with 2 μg/mL puromycin. Human sgRNA sequences: LacZ (CCAGACCGTTCATACAGAAC); NDE1 (g1:GGAACTCCGAGAATTCCAGG, g2: AAAGACTTTCAGCTCCGAGG); KIF2A (GTGGAGATCAAGCGCAGCGA); NEK1 (TAATGGATTACTGTGAGGGA).

#### STAG2/PAXIP1/PAGR1 Knockout Clones

STAG2, PAXIP1, and PAGR1 sgRNAs were cloned into the PX458 backbone and validated with Sanger sequencing. Plasmids were transfected into cells with Lipofectamine 3000 (Invitrogen) per the manufacturer’s guidelines. Seventy-two hours following transfection, the top 10% of GFP+ live cells were sorted into a new population and seeded into a 12-well plate. One week later, the cells were seeded at a concentration of one cell per well in 96-well dishes. Western blots were performed on the clones to confirm complete knockout. Wild-type control cells were generated through transfection of a px458 plasmid containing a LacZ sgRNA. Human sgRNA sequences: LacZ (CCAGACCGTTCATACAGAAC); STAG2 (ATTTCGACATACAAGCACCC); PAXIP1 (AGCCTCACACATAATCTCAG); PAGR1 (TCCCAGACCACACATGCCCA).

#### NDEL1 Heterozygous Knockout Clones

NDEL1 sgRNA (CACCGAGCTAGTTGAATTCCAGGA) was cloned into the PX458 backbone and validated with Sanger sequencing. Knockout and clonal selection were performed as above in the STAG2/PAXIP1/PARGI clones, except that western blots were used to select heterozygous knockout instead of complete knockout.

### shRNA Knockdowns

shRNA sequences corresponding to *NDEL1* or *PAFAH1B1(Lis1)* were cloned into the dox-inducible Tet-pLKO-puro vector (Addgene #21915). Sense and anti-sense oligos for respective shRNA sequences flanked by restriction site overhangs were annealed in a PCR machine with temperature decreasing from 95-20 degrees Celsius. The tet-pLKO-puro vector backbone was digested with the appropriate restriction enzymes and gel-purified. The gel-purified vector and annealed oligos were ligated with T4 DNA ligase (NEB) and transformed into Stbl3 chemically competent cells (ThermoFisher). After growth overnight on ampicillin agar, DNA was extracted from successful colonies and validated by sequencing before transfection and lentiviral transduction into target cells. To induce knockdown, cells were treated with doxycycline at a final concentration of 100ng/ml. Knockdown was confirmed with western blot. shRNA sequences that induced approximately 50% knockdown of each gene were selected for experimental use. Human shRNA target sequences: NDEL1 (GCATCCTTTGATTACTCTCAT, CCGAAAGCTATACCAAATGGT, CCTTCAGTTCAACGACATCTT, CGTCCTTCAGTTCAACGACAT); PAFAH1B1 (GCAGATTATCTTCGTTCAAAT, CGTATGGGATTACAAGAACAA, GCTGAATTAGATGTGAATGAA, TGACCATTAAACTATGGGATT)

### Cell Viability Assays

#### Longitudinal Cell Counting

To evaluate cellular growth dynamics over time, 500,000 cells were seeded in 10cm culture dishes, in triplicate (n=3 plates per condition). When LacZ control plates reached 80% confluency (typically 3-5 days), cells from all conditions were counted using a particle counter (Beckman-Coulter). Counts were recorded, and cells from each condition were re-seeded in triplicate at 500,000 cells per plate. For short-term viability assays, cell counts were taken at first passage. For longitudinal analysis, cells were typically reseeded for a total growth period of 2-3 weeks. Analysis of cell counts was internally normalized to each respective LacZ control within each cell line condition, for example, PC9-LCV2 NDE1 was normalized to PC9-LCV2 LacZ. For visualization, normalized cell counts were multiplied over the duration of the experiment to mimic unrestricted growth.

#### Clonogenic Growth Assays

1,000 to 5,000 cells were plated in triplicate in six-well tissue culture plates directly following genetic knockout and puromycin selection. Cells were allowed to grow in standard conditions for 7-10 days, or until the control reached confluency, to assess the viability effect from the genetic knockout. Each experiment was evaluated against a LacZ control, which was transfected and seeded at the same time as the experimental condition. Plates were then washed with 1X PBS, fixed, and stained with 0.5% (w/v) crystal violet in 6.0% (v/v) glutaraldehyde (Thermo Fisher Scientific). Quantifications of the surface area covered were used to estimate cell viability and were performed using FIJI/ImageJ software with the ColonyArea plugin [59].

### Immunoblotting

Cell pellets were resuspended in 1X RIPA buffer (Sigma) supplemented with Protease and Phosphatase Inhibitor Mini Tablets (Pierce). Cell suspensions were rotated at 4 °C for 15 minutes before 4 °C centrifugation at 13,000×*g* for 10 min. Protein from the resulting supernatant was quantified using the DC Protein Assay (Bio-Rad). The lysates were mixed with Sample Buffer (6X, Bio-Rad) and boiled at 95 °C for 10 min. Samples were run on Bio-Rad 4–20% Bis-Tris Gels and transferred to PVDF membranes using the Trans-Blot Turbo system (Bio-Rad). Membranes were blocked for 1 h with 5% milk/PBS-T (w/v) before incubation with primary antibodies overnight at 4 °C. Primary antibodies were diluted 1:1000 in 5% bovine serum albumin (BSA)/PBS-T (w/v) and were purchased from as follows: β-Actin (CST, #4970), Vinculin (CST, #13901), NDE1 (Proteintech, #10233), NDEL1(Proteintech, #17262), KIF2A (Novus #NB500-180), NEK1, (Proteintech, #27146), Stag2 (#5882). Following primary antibody incubation, membranes were washed with PBS-T and incubated with the corresponding HRP conjugated secondary antibody (CST, #7076s or #7074s) at a dilution of 1:4000 in 5% milk/PBS-t (w/v) for 1 h. Membranes were again washed with PBS-T before developing with Pierce ECL (Thermo) or SuperSignal West Pico chemiluminescent substrate (Thermo) and imaging.

### Immunofluorescence

#### Fixed Imaging

50,000-100,000 lung cancer cells were seeded into 4-well cell culture slides (MatTek) and incubated overnight. After media aspiration, cells were washed once with 1X PBS followed by fixation and permeabilization in 4% paraformaldehyde (v/v), 0.25% Triton X-100 (v/v), and 2 µg/ml Hoechst 33342 stain in PBS for 30 min at room temperature. After dual fixation and permeabilization, cells were washed with 1X PBS and blocked in 3% BSA/PBS for one hour, while protected from light. After aspiration of the BSA, slides were incubated with primary antibody diluted in 3% BSA/PBS (w/v) overnight at 4 °C. Primary antibodies were sourced as follows: cGAS (CST, #15102), Pericentrin (Proteintech, #22271). After primary antibody incubation, cells were again washed with 1X PBS and incubated with corresponding secondary antibodies sourced as follows: Phalloidin-iFlour 633 (Abcam, #AB176758), AlexaFlor 488 goat anti-rabbit IgG (Invitrogen, #A11008) for 1 h at room temperature. Following secondary staining, each well was washed, chamber gaskets were gently removed from each slide, and cover glass was mounted with Dako fluorescence mounting medium (Agilent, S3023). Stained slides were stored protected from light at 4 °C until imaging. Fixed and stained cells were imaged using a 20X objective on a Leica SP5 inverted confocal microscope. A minimum of at least five images from non-overlapping fields of view were taken for each condition. All image analysis was performed using FIJI/ImageJ software. Quantifications were performed by at least two separate individuals using a combination of manual feature identification and established protocols from Cell Profiler 4 [60]. Mitotic index was calculated as the percentage of cells in a given frame in active mitosis (from DNA condensation to daughter cell separation).

#### Live Cell Imaging

To allow visualization of DNA through mitosis, PC9 and PC9-BrM3 cells were stably transfected with FU-H2B-GFP (Addgene, #69550). Within 3-5 days of initiating imaging, cells were transfected with LCV2-sgLacZ or LCV2-sgNDE1 as described above. After selection, approximately 100,000 cells were plated into 35mm glass-bottom petri dishes (MatTek) and incubated overnight to allow for attachment to the culture plate. Live cell imaging was performed on asynchronous cells using the 20X objective on a VivaView FL Incubator Microscope. Brightfield and fluorescent images were captured at 10-minute intervals throughout a 72-hour imaging period, across five distinct fields of view per condition. All live cell imaging analysis was performed using MetaMorph software. Quantifications of multipolarity were performed by manual feature identification by two individuals (n=100 cells/condition). Time in mitosis was calculated as the elapsed duration between DNA condensation and daughter cell separation, as timestamped in the MetaMorph software (n=100 cells/condition).

### DepMap Analyses

#### Correlation Analyses

DepMap correlation analyses were performed in the custom analysis browser using the Pearson correlation analysis, and data were downloaded for visualization in GraphPad. For snake plots, genes were ranked by correlation t-statistic, as generated by the analysis browser in DepMap. CRISPR gene effect sizes are from DepMap, Public 22Q4+score, Chronos. Expression data is from the 22Q2 public release.

#### STAG2/PAXIP1/PAGR1 Signature

To generate the *STAG2*, *PAXIP1*, and *PAGR1* signatures, we correlated the Expression Public 23Q4 DepMap dataset with the expression of either *STAG2*, *PAXIP1*, or *PAGR1*, respectively, in solid tumor cell lines. For each gene, the top 100 correlates by correlation coefficient were included in the gene signature. We then performed GSEAPreranked using GSEA v4.2.3 on the ranked list of differential gene expression analysis from our RNAseq dataset comparing PC9 and PC9-BrM3 cell lines.

### Overexpression Studies

The NDEL1 and LIS1 open reading frames were separately shuttled from pcDNA3 Ndel1 (Addgene #12571) and pcDNA3 Lis1 (Addgene #12574) into N174-MCS (Addgene ##81061) using PCR amplification, restriction digest, and subsequent ligation. Both constructs were sequence validated by Sanger sequencing. Lentivirus was generated from N174-NDEL1 and N174-Lis1 and used to infect cell lines as described above for LCV2 constructs. After selection with 2 μg/mL puromycin, for 48 hours, overexpression was confirmed via western blot.

### Flank Xenograft Studies

Dox-inducible NDE1 knockout constructs were generated by cloning NDE1 or LacZ control sgRNAs into the pRDA_355 backbone as described above for LVC2. Lentivirus was generated for each construct as described above. PC9 or PC9 NDEL1 heterozygous knockout (PC9-NDEL1-C8) cell lines were stably transfected with pKLV2-EF1a-Cas9Bsd-W and kept under blastocidin selection at 10g/ml to maintain Cas9 expression. After Cas9 expression was confirmed, cells were infected with pRDA_355 containing sgRNAs against either LacZ or NDE1 and selected with 2 μg/mL puromycin. Cells expressing pRDA_355 constructs were grown in culture media containing tetracycline-free FBS (VWR). 1 × 10^6^ PC9 or PC9-NDEL1-C8 cells containing the appropriate pRDA_355 constructs were subcutaneously injected into 10-week-old female athymic nu/nu mice flanks in a 1:1 ratio with Matrigel. Once tumors reached 100 mm^3^, they all began doxycycline treatment. Tumors that failed to grow larger than 100mm^3^ in any condition were excluded from further analysis. Doxycycline was prepared fresh at 10mg/ml in water, and administered at 100ul/mouse via oral gavage for 10 days, after which mice were given Doxycycline Chow (DOX rodent chow (625ppm, Rodent Diet (2018, 625 Dox, B)). All tumors were measured via calipers every 1 to 4 days, and tumor volume was calculated *V* = (*L* × W × W)/2 (L = longest diameter and W = shortest diameter). The study continued until IACUC-approved endpoints, including when tumors reached ∼1,500 mm^3^, tumors were ulcerated, or the mice displayed clinical impairments.

## Statistical Analysis

Unless otherwise stated, all results are representative of at least 3 biological replicates, shown as means with SEM used for error visualization. Statistical analyses were performed using GraphPad Prism 9 software. Mouse numbers per group were assessed through power calculations (α = 0.05), where 10 mice per group allows for 90% power to detect inter-group differences of 50% and assuming intra-group variability of 25%. For Kaplan-Meier survival analysis, *p*-values were calculated using log-rank (Mantel-Cox) testing. *P* values were determined using unpaired, two-tailed Student *t* tests, or, for grouped analyses, one-way or two-way ANOVA with the Tukey *post hoc* test. Key: * *p* < 0.05, ** *p* < 0.01, ***, *p* < 0.005, **** *p* < 0.0001.

## Acknowledgments

We thank Drs. Xinna Zhang and Xiongbin Lu (Indiana University) for providing isogenic DU145 prostate cancer cells harboring or lacking heterozygous Chr 17p loss. Similarly, we thank Dr. Beth Sullivan (Duke University) for providing RPE-hTERT cells. We are thankful for the guidance of Dr. Yasheng Gao and the Duke Light Microscopy Core Facility for training and assistance with the microscopy studies. We acknowledge support from NIH grants R01 CA263593 (to K.C.W.), F31 CA287604 (to C.M.T.), and F99/K00 CA245732 (to J.P.H.), DoD Lung Cancer Research Program grants HT9425-24-1-0338 and HT9425-25-1-0714 (to K.C.W.), and V Foundation for Cancer Research grant AST2025-009 (to K.C.W.). T.B.Y is supported by a fellowship from the Edmond J. Safra Center for Bioinformatics at TAU.

## Supplemental Figures

**Fig. S1.**
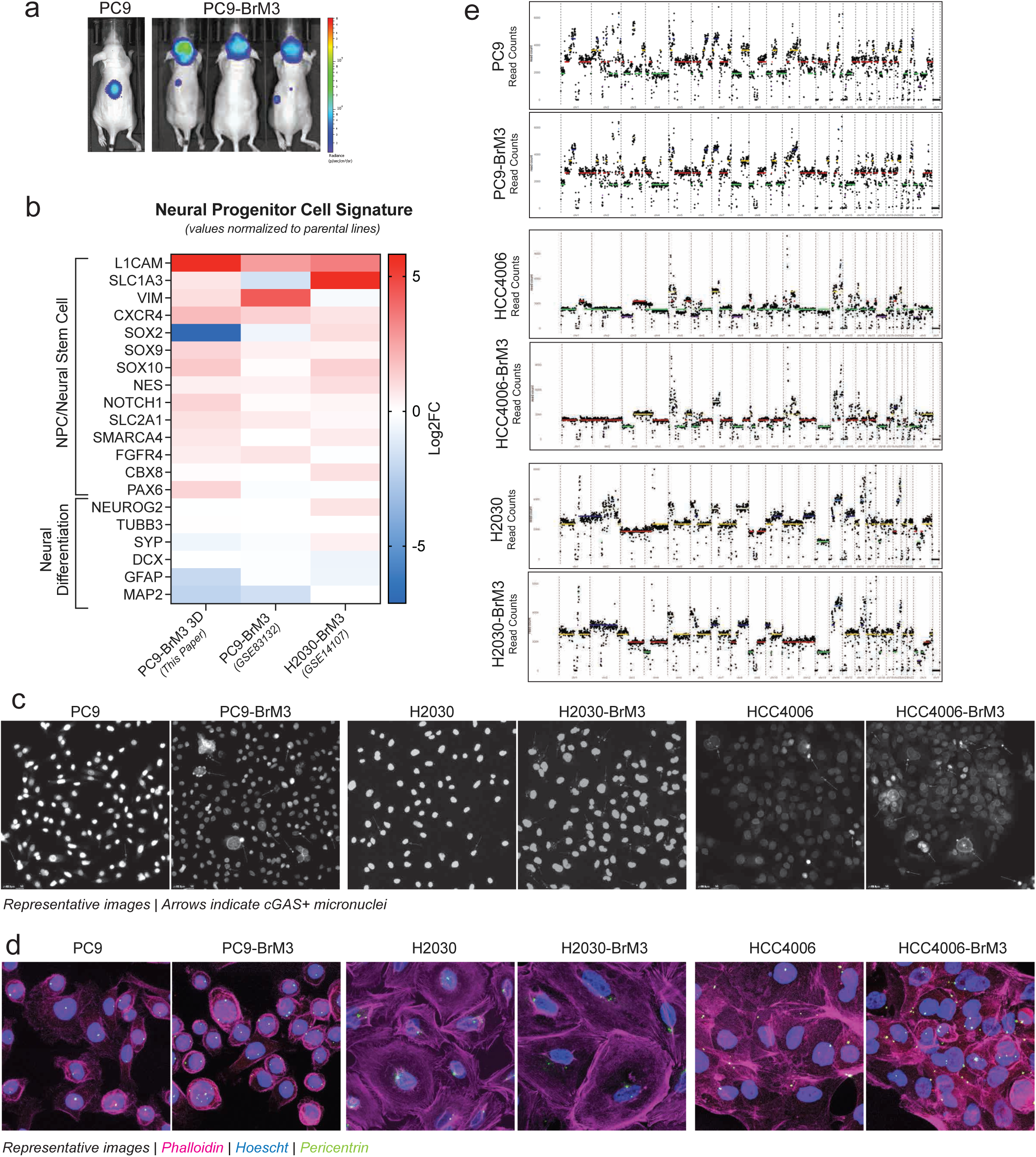
**(A)** Representative bioluminescence imaging of PC9 parental cells and PC9-BrM3 derivative cells after intracardiac injection. **(B)** Heat map of neural stem cell gene expression across BrM derivative cell lines from this study and previously published studies using the same derivation methodology. Values are normalized to respective parental cell line pairs. **(C)** Representative immunofluorescence images from cGAS+ micronuclei quantification in Fig. 1a. **(D)** Representative immunofluorescence images from centrosome quantification in Fig. 1b. **(E)** Low-pass whole genome sequencing read counts generated by AneuFinder [51], showing chromosome arm losses and gains within each cell line and BrM3 derivative.

**Fig S2.**
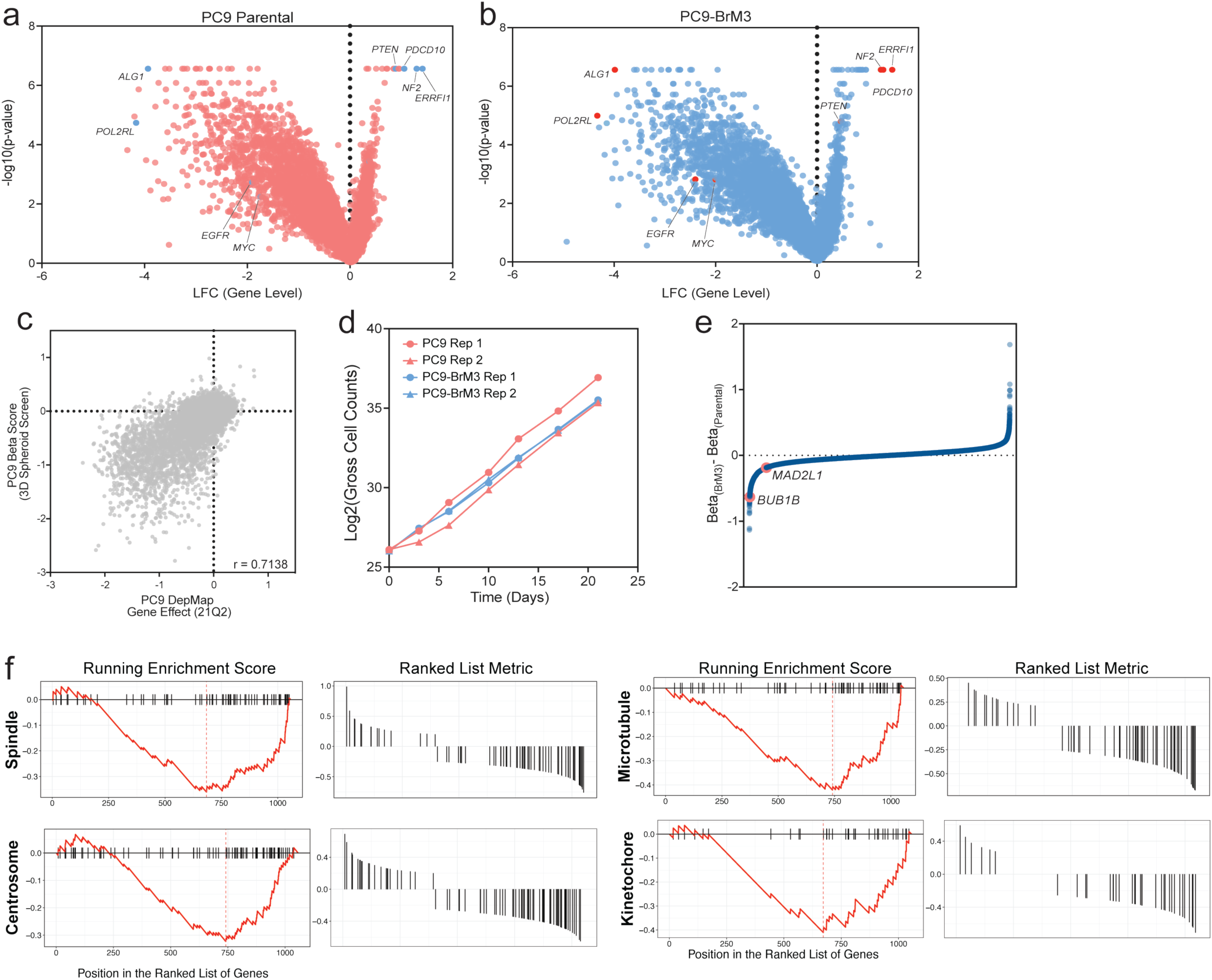
**(A)** Gene-level log fold change of relevant genes from the CRISPR/Cas9 screen in parental PC9 cells. **(B)** As in (A), for PC9-BrM3 cells showing similar dependency profiles for genes known to be either positive or negative regulators of fitness in PC9 cells. **(C)** Scatterplot comparing gene effect scores from the PC9 parental arm of our 3D metastatic screen to 2D gene effect scores from DepMap, showing relatively high degree of similarity between screening modalities. **(D)** Cell counts over the course of the metastatic screen across each replicate in PC9 and PC9-BrM3 cells. **(E)** Snake plot of the beta score difference between PC9 parental and PC9-BrM3 cells, with *BUB1B* and *MAD2L1* highlighted. **(F)** Select running enrichment plots from the GSEA analysis in Fig. 1f.

**Fig S3.**
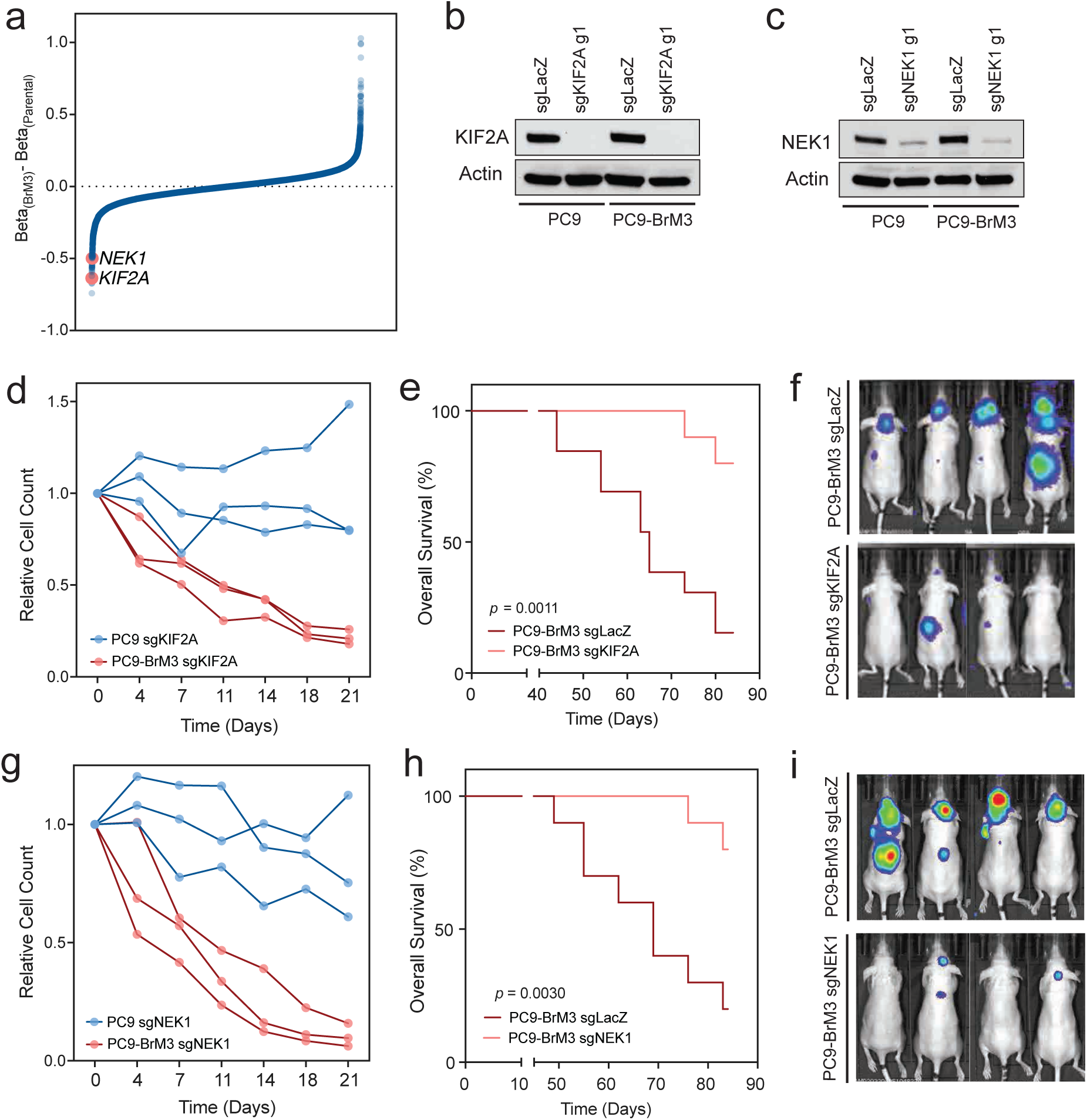
**(A)** Snake plot of the beta score difference between PC9 parental and PC9-BrM3 cells, with *KIF2A* and *NEK1* highlighted. **(B)** Western blot measuring KIF2A levels in PC9 and PC9-BrM3 cells after transfection with LCv2-sgKIF2A and subsequent puromycin selection. **(C)** Immunoblot measuring NEK1 levels in PC9 and PC9-BrM3 cells after transfection with LCv2-sgNEK1 and subsequent puromycin selection. **(D)** Longitudinal cell counting assay comparing cell growth between PC9 cells with *KIF2A* knockout and PC9-BrM3 cells with *KIF2A* knockout, normalized to LacZ targeting sgRNA controls. **(E)** Kaplan-Meier curves of overall survival in mice injected intracardially with PC9-BrM3 cells transduced with Cas9 and sgRNAs against LacZ (control) or *KIF2A*. **(F)** Representative bioluminescent images of mice from (E). **(G-I)** same as (D-F), with *NEK1* knockout.

**Fig. S4.**
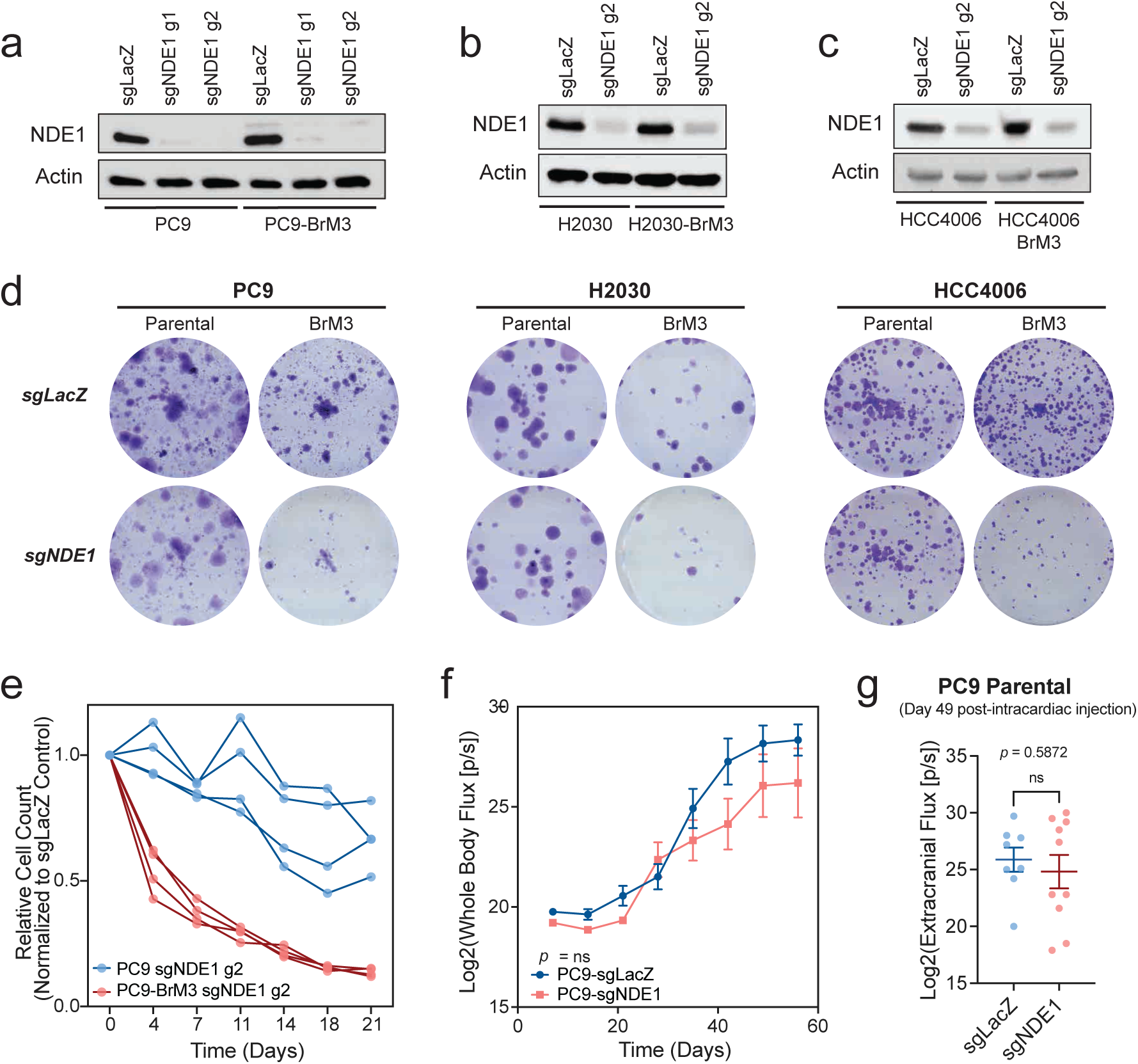
**(A)** Western blot measuring NDE1 levels in PC9 and PC9-BrM3 cells after transfection with LCv2-sgNDE1 and subsequent puromycin selection, across two sgRNAs. **(B)** Western blot measuring NDE1 levels in H2030 and H2030-BrM3 cells after transfection with LCv2-sgNDE1 and subsequent puromycin selection. **(C)** As in (B), with HCC4006 and HCC4006-BrM3 cells. **(D)** Representative images of clonogenic assays from Fig. 2c. **(E)** Longitudinal cell counting assay comparing cell growth between PC9 cells with *NDE1* knockout and PC9-BrM3 cells with *NDE1* knockout, normalized to LacZ targeting sgRNA controls. (Related to Fig. 2b, using a different sgRNA sequence). **(F)** Whole body flux measurements from BLI imaging of mice injected intracardially with PC9 parental cells transduced with Cas9 and sgRNAs against LacZ (control) or *NDE1*. **(G)** Extracranial flux measurements from BLI imaging of mice in (F), on day 49 post-intracardiac injection.

**Fig. S5.**
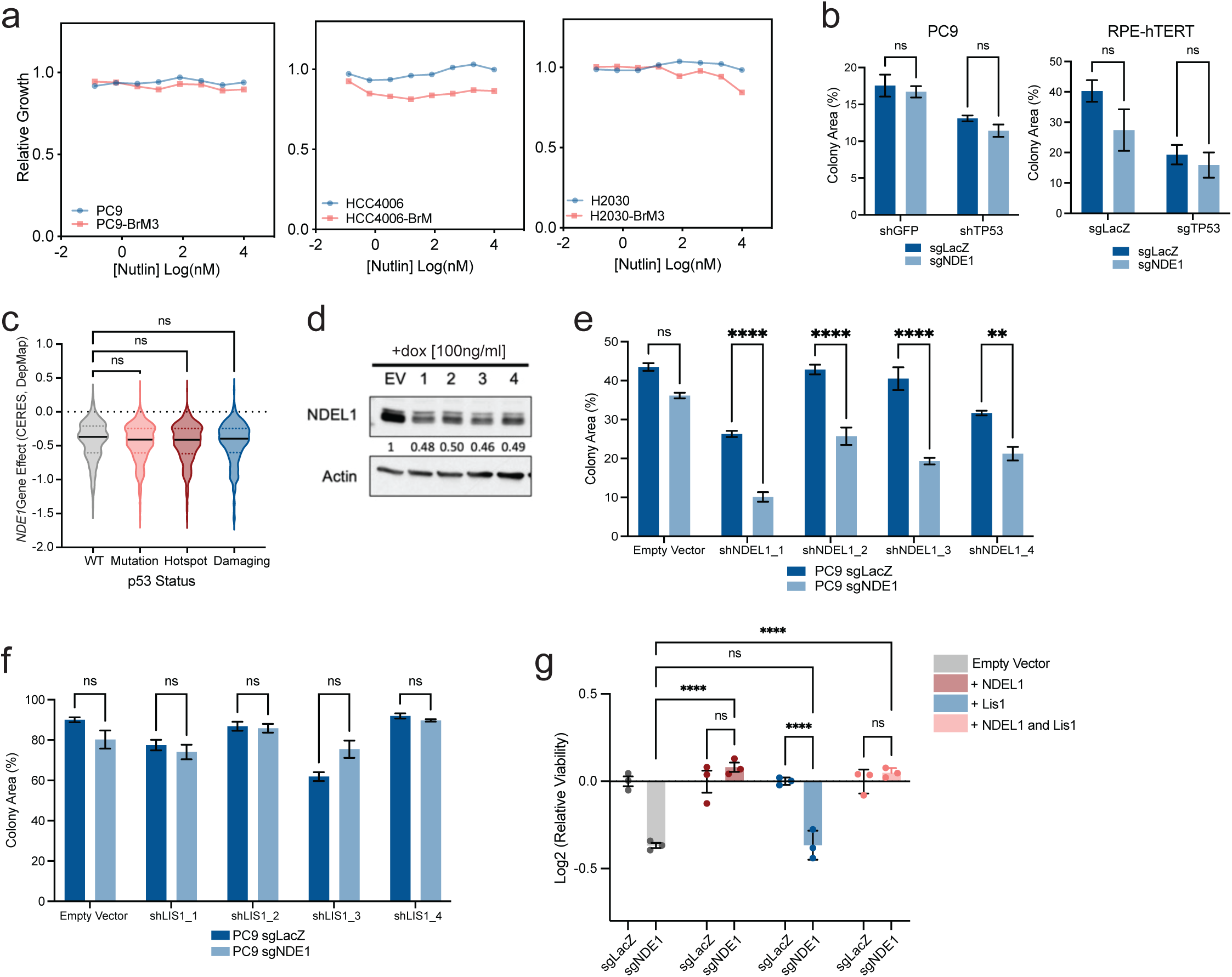
**(A)** IC50 curves of parental and metastatic cell line pairs treated with Nutlin, measured via cell titer glo, normalized internally to DMSO controls. **(B)** Clonogenic growth assays comparing sensitivity to *NDE1* knockout in PC9 and RPE-hTERT cells after depletion of *TP53*. **(C)** Violin plot of *NDE1* gene effect across cell lines in DepMap, segmented by *TP53* mutation status. **(D)** Western blot showing relative knockdown of NDEL1 across 4 selected shRNA constructs and an empty vector (EV) control. Numbers below the NDEL1 blot are relative protein quantifications normalized to EV control, as assessed by blot signal intensity in ImageJ. **(E)** Clonogenic growth assays measuring sensitivity to *NDE1* knockout in PC9 cells after *NDEL1* knockdown with 4 separate shRNAs from (D). **(F)** Clonogenic growth assays measuring sensitivity to *NDE1* knockout in PC9 cells after *PAFAH1B1*/*LIS1* knockdown with 4 separate shRNAs. **(G)** Cell counting assay measuring rescue of *NDE1* dependence in HT-29 cells after overexpression of *NDEL1* and *LIS1*.

